# E2F/Dp inactivation in fat body cells triggers systemic metabolic changes

**DOI:** 10.1101/2021.04.08.439036

**Authors:** Maria Paula Zappia, Ana Guarner, Nadia Kellie-Smith, Alice Rogers, Robert Morris, Brandon Nicolay, Myriam Boukhali, Wilhelm Haas, Nicholas J. Dyson, Maxim V. Frolov

## Abstract

The E2F transcription factors play a critical role in controlling cell fate. In *Drosophila*, the inactivation of E2F in either muscle or fat body results in lethality, suggesting an essential function for E2F in these tissues. However, the cellular and organismal consequences of inactivating E2F in these tissues are not fully understood. Here, we show that the E2F loss exerts both tissue-intrinsic and systemic effects. The proteomic profiling of E2F-deficient muscle and fat body revealed that E2F regulates carbohydrate metabolism, a conclusion further supported by metabolomic profiling. Intriguingly, animals with E2F-deficient fat body had a lower level of circulating trehalose and reduced storage of fat. Strikingly, a sugar supplement was sufficient to restore both trehalose and fat levels, and subsequently, rescued animal lethality. Collectively, our data highlight the unexpected complexity of *E2F* mutant phenotype, which is a result of combining both tissue-specific and systemic changes that contribute to animal development.

## INTRODUCTION

The mechanisms by which E2F transcription factors regulate cell cycle progression have been studied in detail. The activity of E2F1 is restrained by the retinoblastoma protein pRB, a tumor suppressor that is either mutated or functionally inactivated in various cancers (Dyson, 2016). In the textbook description of E2F regulation, the cyclin-dependent kinases phosphorylate pRB, releasing E2F1 to activate the expression of genes that regulate DNA synthesis, S-phase entry and mitosis.

This textbook model, however, is incomplete. Several studies have shown that E2Fs have additional roles that extend beyond the cell cycle. Notably, E2F can also act as a regulator of cellular metabolism (Denechaud et al., 2017; Nicolay and Dyson, 2013). E2F1 was implicated in global glucose homeostasis by controlling insulin secretion in the pancreatic beta cells (Annicotte et al., 2009) and it is needed to promote adipogenesis (Fajas et al., 2002). E2F1 and pRB have been shown to form repressor complexes on the promoters of genes involved in oxidative metabolism and mitochondrial biogenesis (Blanchet et al., 2011), and E2F1 has been found to activate the expression of glycolytic and lipogenic genes in the liver (Denechaud et al., 2016). Accordingly, the inactivation of the RB pathway results in profound metabolic alterations, including changes in central carbon metabolism that confer sensitivity to oxidative stress (Nicolay et al., 2013; Reynolds et al., 2014).

The model organism *Drosophila* has provided important insights into our current understanding of E2F control. The mechanisms of action of pRB and E2F orthologues are highly conserved from flies to mammals (Van Den Heuvel and Dyson, 2008). The *Drosophila* E2F family contains two E2F genes, *E2f1* and *E2f2*. Each E2F forms a heterodimer with the binding partner Dp, which is required for high-affinity DNA binding. An important feature of the Drosophila E2F/RB network is that Dp is encoded by a single gene. As a result, the entire program of E2F regulation can be abolished by the inactivation of Dp, either through a *Dp* mutation or in a tissue-specific manner by RNA interference (RNAi) (Frolov et al., 2005; Guarner et al., 2017; Royzman et al., 1997; Zappia and Frolov, 2016). Studies of *Dp* loss of function provide a glimpse of the overall function of E2F; a perspective that has not yet been possible in studies in mammalian cells that have far larger families of E2F, DP and RB proteins.

The loss of E2F function, as seen in *Dp* mutants, is permissive for development until pupation when lethality occurs. This confirmed the general expectation that E2F/DP will be absolutely essential for animal viability. Surprisingly, hallmarks of E2F regulation, such as cell proliferation, differentiation or apoptosis were largely unaffected in *Dp* mutants (Frolov et al., 2001; Royzman et al., 1997). Interestingly, the inactivation of E2F in either the fat body, which serve the roles of liver and adipose tissue, or muscles phenocopied the lethal phenotype of *Dp* mutants pointing to the requirement of E2F for animal viability in both tissues (Guarner et al., 2017; Zappia and Frolov, 2016).

The lethality of *Dp* mutants can be rescued by tissue-specific expression of *Dp* in either the muscles or fat body (Guarner et al., 2017; Zappia and Frolov, 2016). These findings confirm the importance of Dp in these tissues but, taken together, the results are puzzling: it is not known why *Dp* expression in one of these tissues can rescue an essential function provided in the other. Answering this question is complicated by the fact that the cellular consequences of *Dp* loss in the muscles or fat body are not understood in detail. Given the inter-communication between muscle and fat body (Demontis et al., 2014; Demontis and Perrimon, 2009; Zhao and Karpac, 2017), it was possible that E2F inactivation in muscle might affect the fat body, and/or *vice versa*. A further possibility was that defects resulting from tissue-specific *Dp* depletion might generate systemic changes.

To address these questions, and to ask whether the roles of E2F in muscle and fat body are the same or different, we examined the changes that occur when *Dp* is specifically removed from each of these tissues. Quantitative proteomics and metabolomic profiling of both *Dp*-deficient tissues revealed changes in carbohydrate metabolism. *Dp*-deficiency in the fat body resulted in low levels of trehalose, the main circulating sugar in hemolymph, and abnormal triglycerides storage. Strikingly, these defects were suppressed on high sugar diet that also rescued the lethality of *Dp*-deficient fat body animals. Despite finding that *Dp-* deficient fat bodies and muscles share similar proteomic changes in carbohydrate metabolism, rearing larvae on high sugar diet had no beneficial effect on animals that lacked Dp function in muscles. Taken together these findings show that E2F has important metabolic functions in both muscles and fat body and that the loss of this regulation, at least in the fat body, leads to systemic changes that can be suppressed by a high sugar diet. These observations show that E2F regulation is needed to prevent both tissue specific and systemic phenotypes.

## RESULTS

### The loss of E2F/DP in the fat body does not impact muscle development

The expression of *UAS-Dp* RNAi transgene driven by the muscle (*Mef2-GAL4*) or fat body (*cg-GAL4*) specific GAL4 drivers specifically depletes Dp protein in the corresponding tissue and results in readily observed phenotypes: the inactivation of Dp in muscle is accompanied by severely reduced muscle growth (Figure 1A-C) (Zappia and Frolov, 2016), while binucleated cells and decreased fat storage were found following fat body-specific Dp depletion (Figure 2A) (Guarner et al., 2017). Given that the inactivation of Dp in either tissue had a similar impact on viability and caused pupal lethality (Figure S1) we asked whether Dp depletion in muscle may also affect fat body development and, conversely, if Dp deficiency in fat body may cause muscle abnormalities.

**FIGURE 1:**
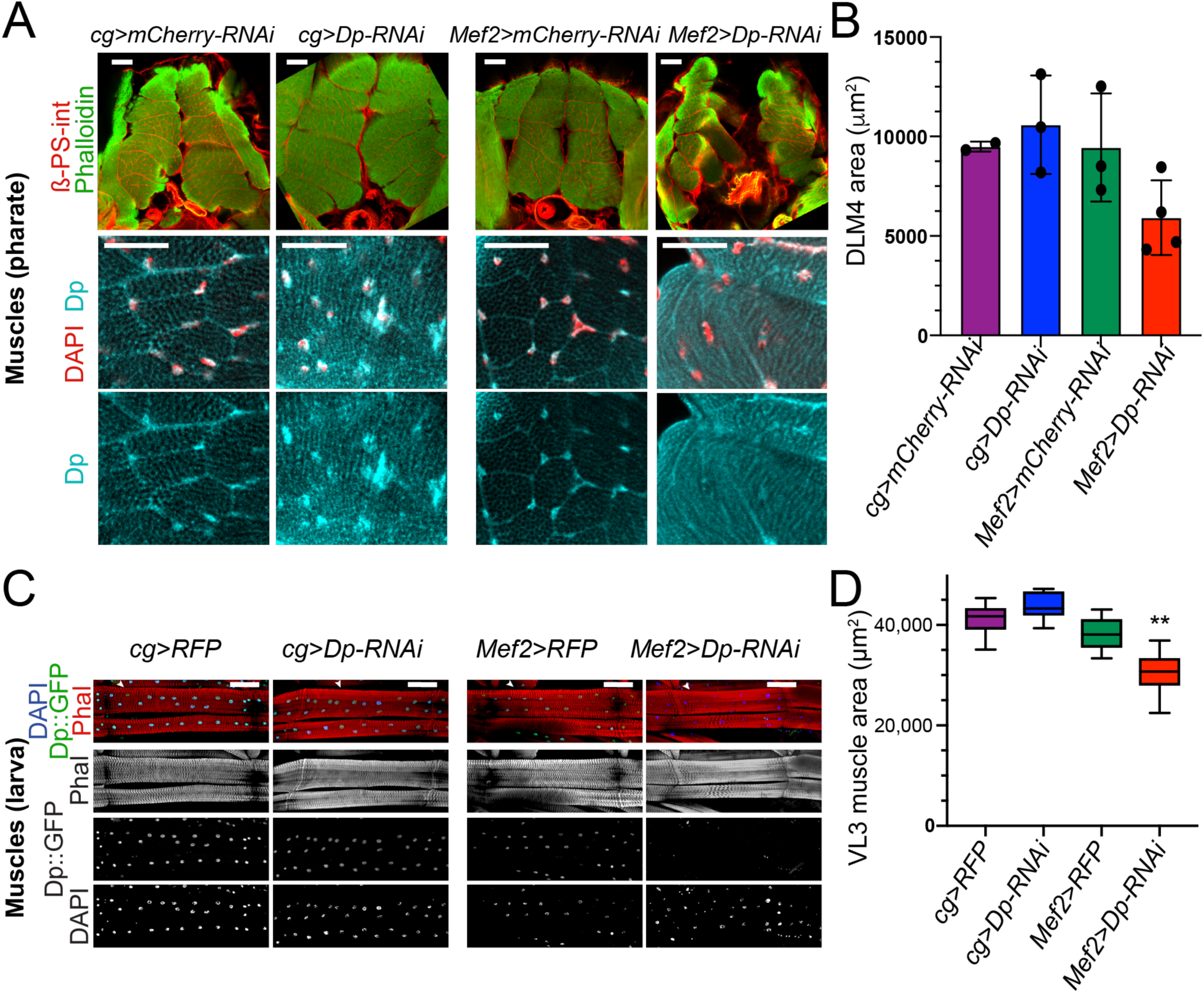
The loss of E2F in fat body does not impair muscle formation. **(A)** Confocal single section images of transverse sections of thoraces at 96 h APF stained using anti-ß-PS-integrin antibody, phalloidin, anti-Dp antibodies (212) and DAPI. Ventral to bottom. Magnification of DLM4 is in bottom panel. Scale: 50 μm. **(B)** Quantification of DLM4 area. Data are represented as mean ± SD. Kruskal-Wallis test, p=0.09. n=2-4 animals per genotype. **(C)** Confocal Z-stack-projected images of third instar larval body wall muscles VL3 (marked with white arrowhead) and VL4 from the segment A4 stained with Phalloidin, Dp::GFP and DAPI. Anterior is to the left. Scale: 100 μm. **(D)** Box plot showing quantification of VL3 muscle area. Whiskers are min to max values, Kruskal-Wallis test followed by Dunn’s multiple comparisons test, ** p<0.0001. n=10-12 animals per genotype. Three independent experiments were done. Full genotypes are (A-B) *cg-GAL4,UAS-mCherry-RNAi*, *cg-GAL4/UAS-Dp[GD4444]-RNAi*, *Mef2-GAL4/UAS-mCherry-RNAi*, and *UAS-Dp[GD4444]-RNAi,Mef2-GAL4* (C-D) *cg-GAL4/UAS-RFP,Dp[GFP]*, *cg-GAL4/UAS-Dp[GFP],UAS-Dp[GD4444]-RNAi*, *UAS-RFP,Dp[GFP];Mef2-GAL4*, and *UAS-Dp[GFP],UAS-Dp[GD4444]-RNAi,Mef2-GAL4*.

**FIGURE 2:**
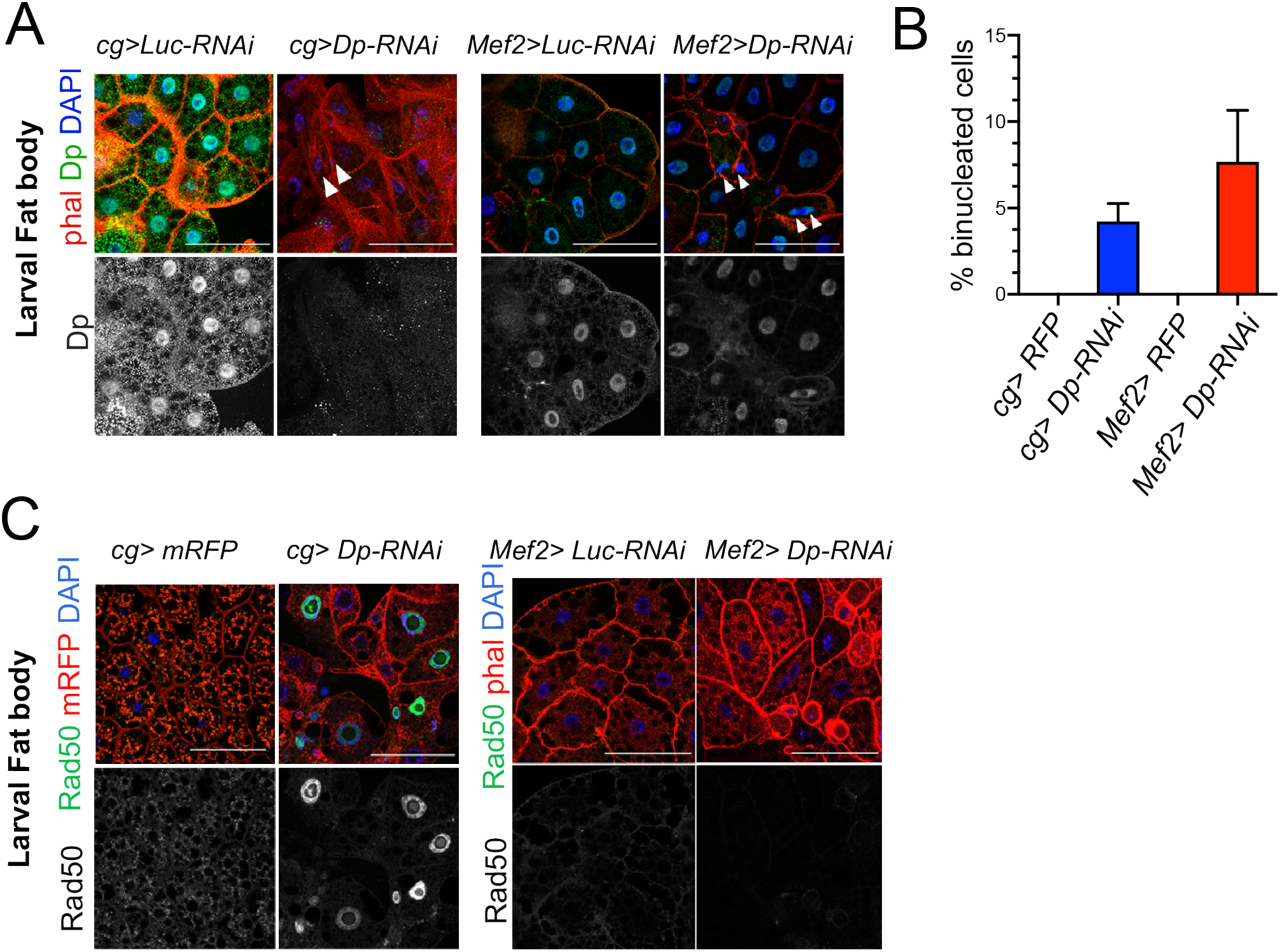
The loss of E2F in muscles has a systemic effect in the development of fat body. **(A)** Confocal single plane images of third instar larval fat bodies stained with phalloidin, DAPI and mouse anti-Dp antibody (Yun). White arrowheads point to newly formed binucleate cells. **(B)** Quantification of the percentage of binucleated cells in fat body. Date are presented as Mean ± SD, n=415 cells for *Mef2>Dp-RNAi*, n=405 cells for *cg>Dp-RNAi*, and n=300 cells for each *Mef2>Luc-RNAi* and *cg>Luc-RNAi*, which did not show binucleates. Kruskal-Wallis test followed by Dunn’s multiple comparisons test, p<0.0001. Experiment was repeated two times. One representative experiment is shown. **(C)** Confocal single plane images of third instar larval fat bodies stained with Rad50, phalloidin and DAPI. Scale: 100 μm. Full genotypes are (A) *cg-GAL4,UAS-luciferase[JF01355]-RNAi*, *cg-GAL4/UAS-Dp[GD4444]-RNAi*, *Mef2-GAL4/ UAS-luciferase[JF01355]-RNAi*, and *UAS-Dp[GD4444]-RNAi,Mef2-GAL4*, (B) *cg-GAL4/UAS-RFP,Dp[GFP], cg-GAL4/UAS-Dp[GFP],UAS-Dp [GD4444]-RNAi*, *UAS-RFP,Dp[GFP];Mef2-GAL4*, and *UAS-Dp[GFP],UAS-Dp[GD4444]-RNAi,Mef2-GAL4*.

**SUPPLEMENTARY FIGURE S1:**
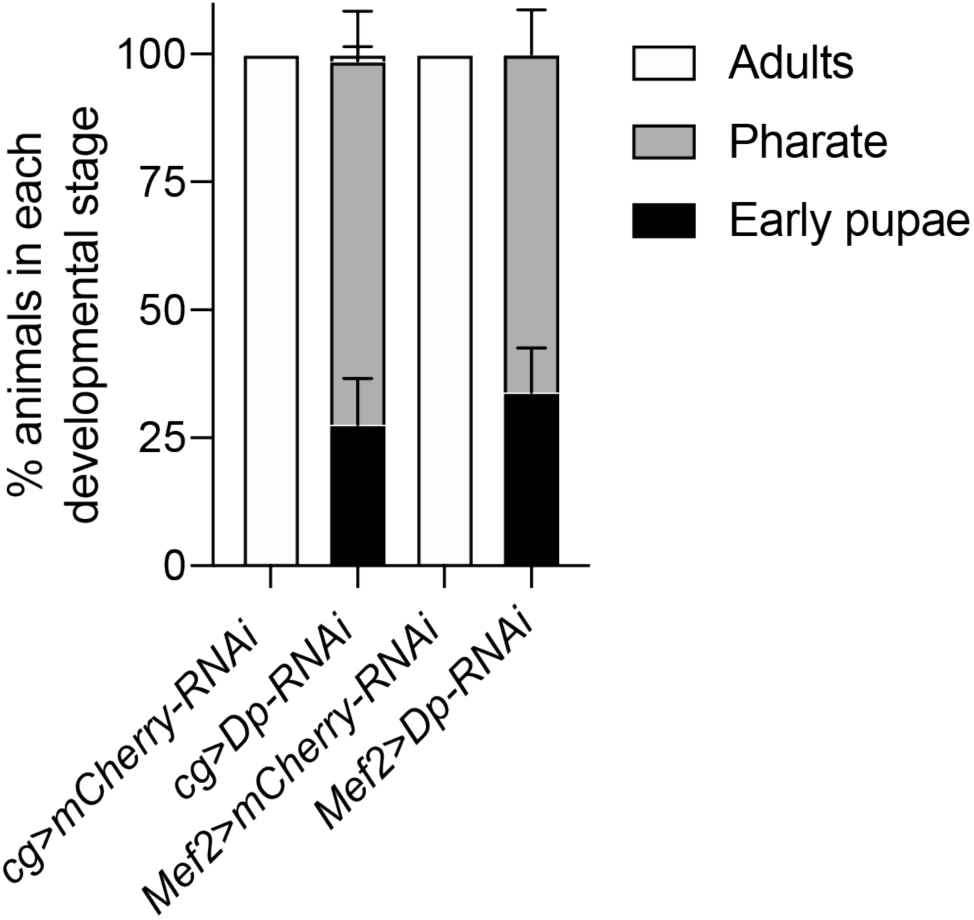
E2F in muscles and fat body is required for animal development. Animal viability assessed by quantifying the percentage of animals at each developmental stage (adult, pharate and early pupa). Data are presented as stacked bar plot, Mean ± SEM, n=2 replicates per genotype, repeated as N=5-6 independent experiments. Full genotypes are *cg-GAL4,UAS-mCherry-RNAi*, *cg-GAL4/UAS-Dp[GD4444]-RNAi*, *Mef2-GAL4/UAS-mCherry-RNAi*, and *UAS-Dp[GD4444]-RNAi*,*Mef2-GAL4*.

We began by comparing muscle structure between *Mef2>Dp-RNAi* and *cg>Dp*-*RNAi* animals. Animals were staged as pharate adults, and cross-sections of thoraces were stained with anti-β-PS integrin antibody and phalloidin to visualize dorsal longitudinal muscles (DLM), the largest muscles in *Drosophila*. As previously reported, DLM of *Mef2>Dp-RNAi* were severely reduced in size (Figure 1A-B) (Zappia and Frolov, 2016). In contrast, no gross differences in muscle morphology was observed in *cg>Dp*-*RNAi* (Figure 1A-B). The efficiency of Dp knockdown was confirmed by staining with anti-Dp antibody in *Mef2>Dp-RNAi* muscles that showed decrease in Dp signal in nuclei (Figure 1A, bottom panel).

The analysis was extended to larval muscles that are also affected by the loss of *Dp* (Zappia and Frolov, 2016). The ventral longitudinal 3 (VL3) muscles in wandering third instar larva were visualized by staining the body walls with phalloidin and DAPI. As previously reported, the area of the VL3 muscles in *Mef2>Dp*-*RNAi* animals were significantly smaller than control. In contrast, no differences were detected in the VL3 muscle area between *cg>RFP* and *cg>Dp*-*RNAi* larval muscles (Figure 1C-D). To confirm efficiency of Dp depletion in *Mef2>Dp*-*RNAi* the expression of Dp was monitored with the transgene *Dp^GFP^*, which expresses the fusion protein *Dp::GFP* (Figure 1C) (Zappia and Frolov, 2016). Thus, we concluded that the loss of E2F/DP in fat body does not induce systemic effects in the developing muscles and does not cause the phenotype observed in Dp-depleted muscles.

### Dp-depleted muscles affect fat body development

One of hallmarks of the loss of *Dp* in fat body is the appearance of binucleated cells (Guarner et al., 2017). We confirmed that the depletion of Dp driven by the fat body*-*specific GAL4 driver *cg-GAL4* resulted in the formation of binucleated cells (∼4.2% in *cg>Dp*-*RNAi*, Figure 2A-B) while none were found in fat bodies from wild type animals. This phenotype was also observed in *E2f2*; *E2f1* double mutant animals (Figure S2A) and in *Dp* null mutants (Guarner et al., 2017). Next, we examined the fat body in *Mef2>Dp-RNAi* larva following muscle-specific Dp depletion by staining the tissue with phalloidin and DAPI. Surprisingly, ablation of Dp in muscles led to the appearance of binucleated cells (7.7% in *Mef2>Dp-RNAi*, Figure 2A-B). We confirmed that Dp was depleted in a tissue-specific manner, as only *cg>Dp*-*RNAi* fat bodies showed reduced levels of Dp protein as revealed by immunofluorescence using anti-Dp antibodies (Figure 2A, bottom panel).

**SUPPLEMENTARY FIGURE S2:**
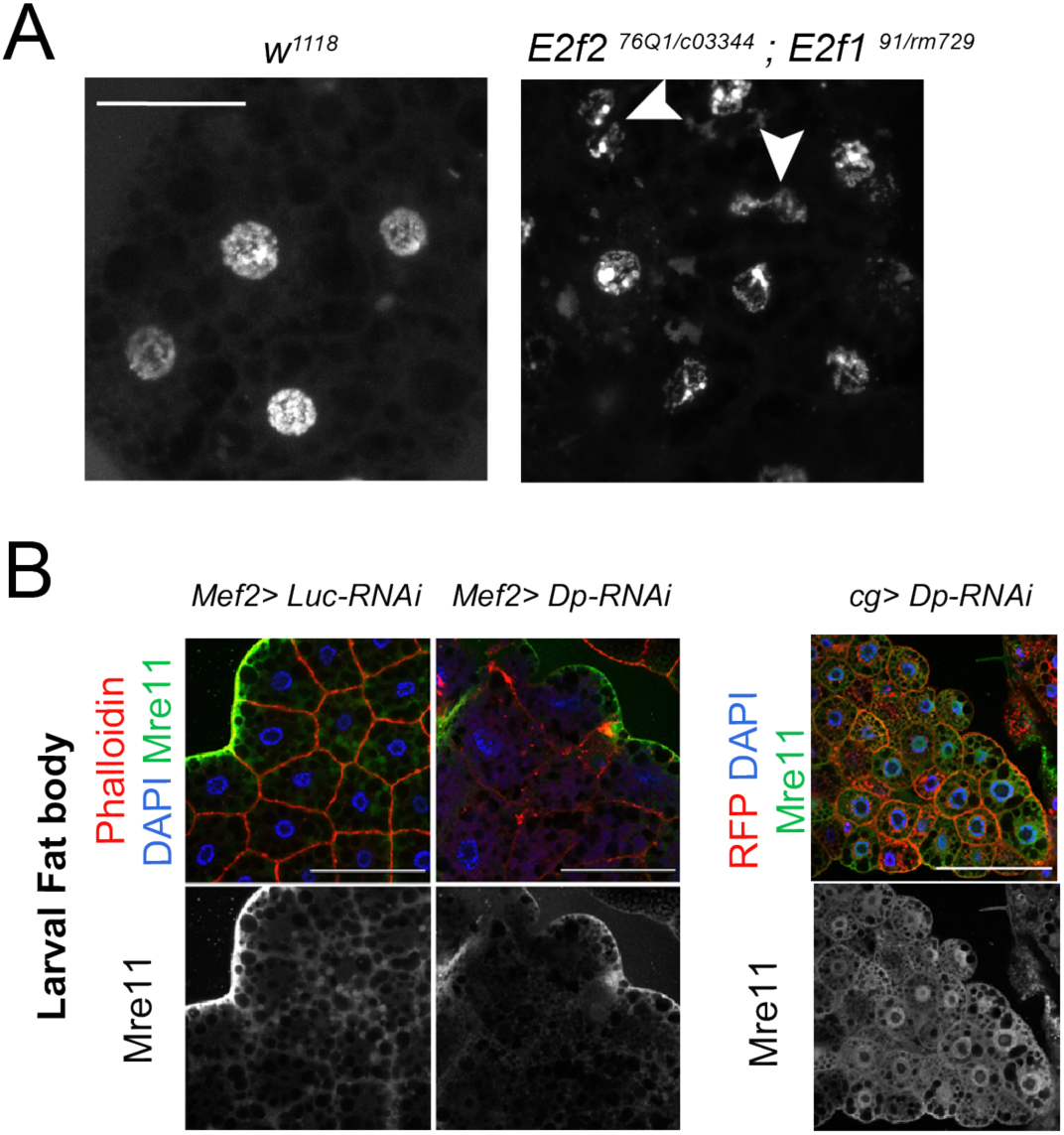
Loss of E2F induces binucleated cells in fat body. **(A)** Confocal single plane images of third instar larval fat bodies stained with DAPI. Genotypes are *E2f2 ^76Q1 /c03344^; E2f1 ^91 / m729^* compared to control *w^1118^*. White arrowheads point to binucleated cells. Scale 50 μm. **(B)** Confocal single plane images of third instar larval fat bodies from *Mef2>Luc-RNAi*, *Mef2>Dp-RNAi* and *cg>DpRNAi* larvae stained with anti-Mre11 antibodies, phalloidin and DAPI. Scale: 100 μm

One of functions of E2F in the fat body is to limit the response to DNA damage. In Dp-depleted fat body, there is an increased recruitment of the DNA damage proteins of the MRN sensor complex, such as Rad50 and Mre11 (Guarner et al., 2017). To determine whether the defects found in the fat bodies of *Mef2>Dp-RNAi* animals were related to the activation of the DNA damage response, tissues were immunostained with anti-Rad50 (Figure 2C) and anti-Mre11 (Figure S2B) antibodies. Notably, the MRN proteins were not recruited in the fat body of *Mef2>Dp*-*RNAi* animals, as opposed to *cg>Dp*-*RNAi* fat body cells.

We conclude that the loss of Dp in muscles elicits defects in fat body that are similar to the phenotype seen in Dp-deficient fat body albeit not accompanied by the upregulation of MRN proteins. Thus, Dp-deficient muscle exerts a systemic effect on normal tissues, such as fat body.

### The loss of Dp in muscles does not alter Dp expression in the fat body

Leaky expression of *GAL4* drivers in other tissues during earlier developmental stages might provide a trivial explanation for the results described above. To exclude this possibility, we used three approaches to confirm the tissue-specificity of *cg-GAL4* and *Mef2-GAL4* drivers used to knockdown Dp. First, we examined the real-time and lineage tracing expression of the drivers. We used the system G-TRACE, which combines the FLP recombinase-FRT and the expression of GFP protein to trace earlier GAL4 expression, and the presence of RFP protein to identify real-time expression of GAL4 (Evans et al., 2009). G-TRACE showed that the *cg-GAL4* driver is expressed in the fat body in agreement with previous report (Pastor-Pareja and Xu, 2011) (left panel, Figure 3A). No GFP or RFP signal was detected in larval muscles (right panel, Figure 3A) suggesting that at no point during development *cg-GAL4* was expressed in muscles. Similarly, *Mef2-GAL4* expression was detected in larval skeletal and smooth muscles, and in the adult muscle precursors (wing disc myoblasts), but not in the fat body (Figure 3B).

**FIGURE 3:**
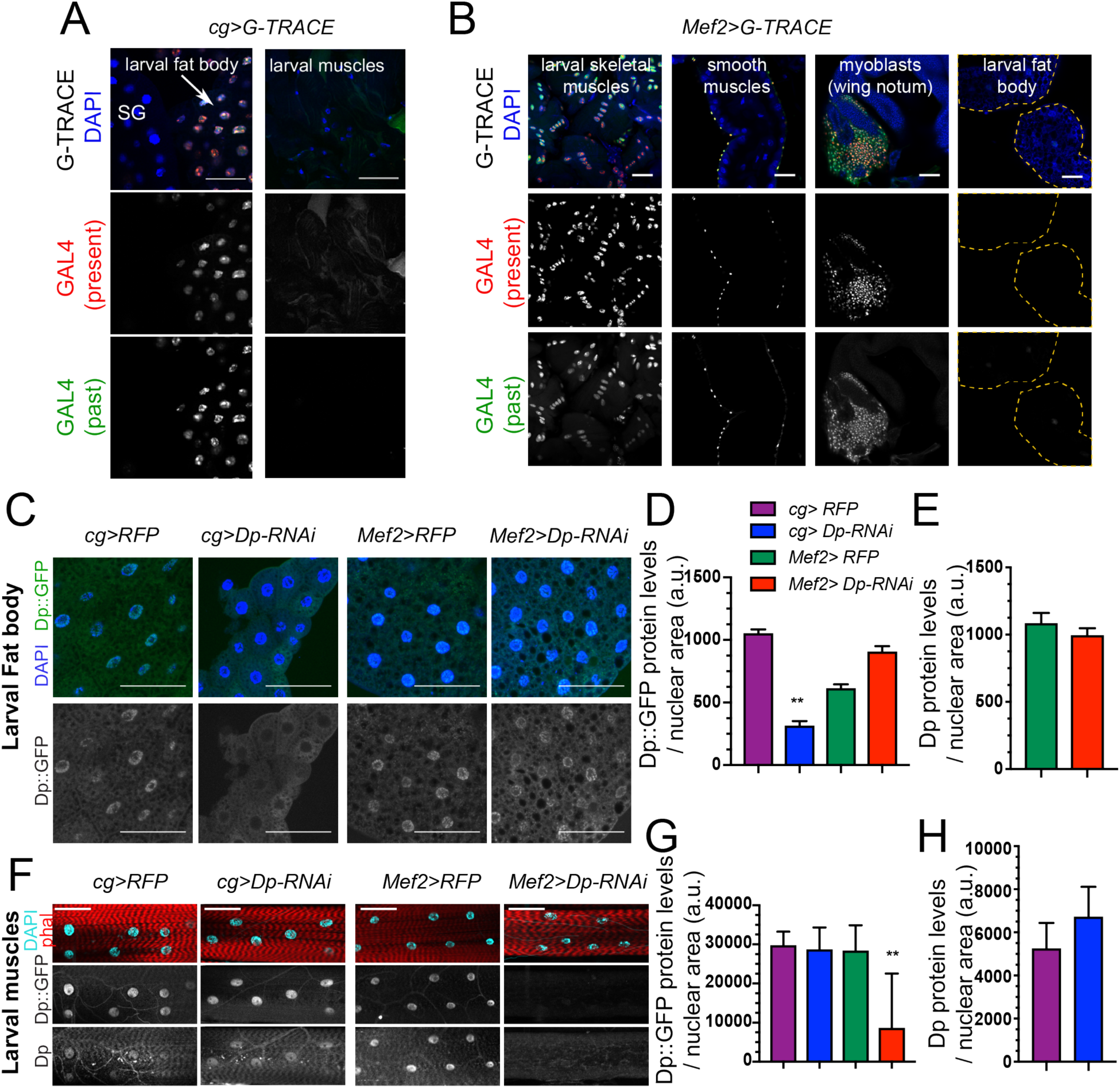
Dp expression is knock-down in a tissue-specific manner. **(A-B)** Lineage tracing of tissues dissected from third instar larvae stained with DAPI and showing the lineage of cg-GAL4 (GFP) and the active GAL4 (RFP). **(A)** Confocal single plane images of *cg>G-TRACE* fat bodies, salivary glands, and muscles. Scale: 100 μm. **(B)** Confocal single plane images of *Mef2>G-TRACE* larval skeletal (body wall) and smooth (gut) muscles, adult myoblasts on the wing discs and fat bodies. Scale: 50 μm. **(C)** Confocal single plane images of third instar larval fat bodies stained with DAPI and showing Dp::GFP tagged protein. White arrowheads indicate binucleated cells. Scale: 100μm. **(D)** Quantification of Dp::GFP protein levels as shown in C, relative to nuclear area. Mean±SEM, Kruskal-Wallis test followed by Dunn’s multiple comparisons test, ** p<0.0001, n=11-20 per genotype. **(E)** Quantification of Dp protein levels relative to nuclear area in larval fat body of *Mef2>RFP* and *Mef>Dp-RNAi* animals. Mean± SEM, Mann-Whitney test, p = 0.35, n=14 per genotype. **(F)** Confocal single plane images of third instar larval muscles immunostained with rabbit anti-Dp antibody (212), phalloidin and DAPI. Scale: 50 μm. **(G)** Quantification of Dp::GFP protein levels as shown in F, relative to nuclear area. Mean± SD, Kruskal-Wallis test followed by Dunn’s multiple comparisons test, ** p = 0.0008, n=10-12 animals per genotype. **(H)** Quantification of Dp protein levels relative to nuclear area in larval fat body of cg>RFP and *cg>Dp-RNAi* animals. Mean ± SD, Mann-Whitney test, p = 0.07, n=6-9 per genotype. Full genotype: (A) cg*-GAL4/UAS-gTRACE* (B) *UAS-gTRACE,Mef2-GAL4*, (C-H) *cg-GAL4/UASRFP, Dp[GFP], cg-GAL4/UAS-Dp[GFP],UAS-Dp [GD4444]-RNAi*, *UAS-RFP,Dp[GFP];Mef2-GAL4*, and *UAS-Dp[GFP],UAS-Dp[GD4444]-RNAi,Mef2-GAL4*.

Second, the levels of Dp expression in fat body and muscles of *Mef2>Dp-RNAi* and *cg>Dp-RNAi* larva were examined by immunofluorescence using Dp antibodies. Third, Dp expression was monitored using a *Dp^GFP^* transgene that expresses a Dp::GFP fusion protein from the endogenous *Dp* locus (Zappia and Frolov, 2016). The Dp::GFP protein was efficiently depleted in fat body of *cg>Dp-RNAi* compared to control (Figure 3C, quantified in Figure 3D). Importantly, the levels of the Dp::GFP protein remained unaffected in the fat body of *Mef2>Dp-RNAi* (Figure 3C, quantified in Figure 3D). This result was further confirmed by staining with anti-Dp antibody that showed no changes in the endogenous expression of Dp in fat body of *Mef2>Dp*-*RNA*i (Figure 3E). Similarly, the expression of Dp::GFP in muscles was not altered in *cg>Dp*-*RNAi*, whereas, as expected, Dp::GFP was significantly reduced in muscles of *Mef2>Dp*-*RNAi* compared to control (Figure 3F, quantified in Fig 3G). Using anti-Dp antibody, we further confirmed that the endogenous levels of Dp protein in muscles did not change upon Dp depletion in the fat body (Figure 3F, quantified in Figure 3H).

Thus, the occurrence of binucleated cells in *Mef2>Dp*-*RNAi* fat body is not due to altered expression of Dp in fat body of these animals and, therefore, reflects a systemic effect induced by muscle-specific Dp depletion.

### The muscle-specific expression of Dp in *Dp* mutants rescues the fat body phenotype

The expression of *UAS*-*Dp* transgene with either the fat body- or muscle-specific drivers, *cg-GAL4* or *Mef2-GAL4*, can significantly extend viability of *Dp* mutants (Guarner et al., 2017; Zappia and Frolov, 2016). Given the systemic effect of Dp described above we asked whether the muscle-specific *Dp* expression suppresses the defects in fat body of *Dp* mutants. The *UAS-Dp* transgene was expressed in the trans-heterozygous *Dp^a3^/Df(2R)Exel7124* (*Dp-/-*) mutant animals under the control of *cg-GAL4* or *Mef2-GALl4*. Larval fat bodies were stained with phalloidin and DAPI to visualize the occurrence of binucleated cells. In agreement with previously published data (Guarner et al., 2017), 7.1% of cells in fat body of *Dp* mutants were binucleated and this phenotype was fully rescued in the *Dp-/-*; *cg>Dp* animals (Figure 4A, quantified in Figure 4B). Strikingly, the binucleated phenotype was also largely rescued by re-expression of *Dp* in muscles of *Dp* mutants, in *Dp-/-*; *Mef2>Dp* animals (Figure 4A-B). We note however, that fat bodies of *Dp-/-*; *Mef2>Dp* animals still contained fragmented and decondensed/large nuclei indicating that the rescue was incomplete. Staining with anti-Dp antibody confirmed the lack of Dp expression in *Dp-/-*; *Mef2>Dp* fat bodies (bottom panel, Figure 4A), thus excluding the possibility of a leaky expression of *Mef2-GAL4* driver in the fat body.

**FIGURE 4:**
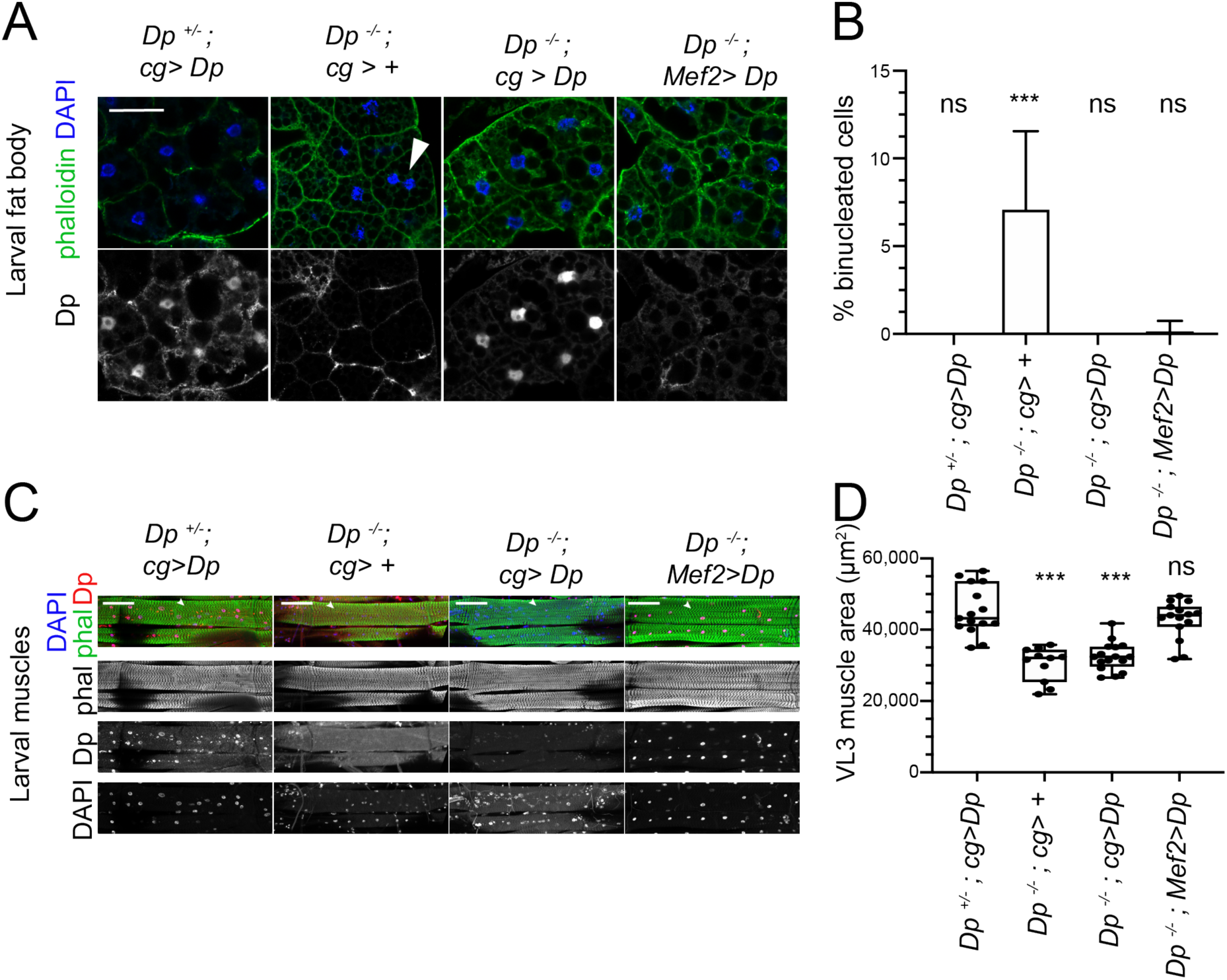
Restoring E2F/Dp in muscles suppresses defect in *Dp*-deficient fat body. **A:** Confocal single plane images of third instar larval fat bodies immunostained with anti-Dp antibody (212), phalloidin and DAPI. Note that *Dp^-/-^;Mef2>Dp* rescued animals do not show binucleated cells, but many nuclei are fragmented and decondensed. Scale: 50 μm **B**: Quantification of percentage of binucleated cells as in A. Data presented as bar plot showing mean ± SD, Kruskal-Wallis test followed by Dunn’s multiple comparisons test, *** p <0.0001, n=16 animals per genotype, two independent experiments were done. At least 606 cells were scored. **C**: Confocal Z-stack-projected images of third instar larval body wall muscles VL3 (marked with white arrowhead) and VL4 from the segment A4 immunostained with rabbit anti-Dp antibody (212), phalloidin and DAPI. Anterior is to the left. Scale: 100 μm **D**: Quantification of VL3 muscle area as in C. Data presented as box plot, whiskers min to max values, Kruskal-Wallis test followed by Dunn’s multiple comparisons test, *** p <0.0001, n=15 animals per genotype, except n=10 for Dp-/-, two independent experiments were done. Full genotypes are *Dp^+^/Dp^a3^,cg-GAL4;UAS-Dp, Dp^Exel7124^/Dp^a3^,cg-GAL4*, *Dp^Exel7124^/Dp^a3^,cg-GAL4;UAS-Dp*, *Dp^Exel7124^/Dp^a3^; Mef2-GAL4/UAS-Dp*

Next, we asked whether re-expression of *Dp* in the fat body suppresses muscle defects of *Dp* mutants. The body walls of third instar larvae were dissected and VL3 muscles of the A4 segment were visualized by staining the tissue with DAPI and phalloidin. As previously reported, VL3 muscle area was smaller in *Dp*-/-mutant larvae and fully rescued in *Dp-/-*; *Mef2>Dp* (Figure 4C, quantified in Figure 4D) (Zappia and Frolov, 2016). In contrast, the expression of *Dp* in the fat body was insufficient to suppress the small size of VL3 muscles in *Dp-/-* mutant larvae (Figure 4 C, quantified in Figure 4D).

Thus, muscle-specific expression of *Dp* can rescue the binucleated phenotype of *Dp* mutant fat body, which is consistent with the idea that E2F/Dp in muscle exerts a systemic effect in larva that impacts the fat body. In contrast, re-expressing *Dp* in the fat body is insufficient to suppress the muscle defects in *Dp* mutants. This suggests that E2F/Dp modifies the inter-tissue communication between muscle and fat body.

### Integrating proteomic and metabolomic profiling of E2F-depleted tissues uncovers alterations in carbohydrate metabolism

A significant limitation of transcriptional profiles is that it is difficult to know whether a change in mRNA levels leads to a measurable difference in protein level, or causes a change in pathway activity. To avoid these issues, we generated proteomic profiles of fat body and skeletal muscles and used these to compare how the loss of E2F affects protein levels in each tissue. Third instar larval fat bodies were collected from both wild type (*w^1118^*) and *Dp-/-* mutant (Figure 5A, left panel) and subjected to multiplexed quantitative mass spectrometry-based proteomics using tandem-mass tag (TMT) technology (Edwards and Haas, 2016; McAlister et al., 2014). Collecting larval muscles in sufficient quantities for such proteomic profiling was not feasible due to technical challenge of separating larval muscle from adjacent tissue. Therefore, we turned to dissecting thoracic muscles from pharate pupa. We used *Mef2>Dp*-*RNAi* pharates since *Dp-/-* mutants die as early pupa (Figure 5A, right panel). Western blot analysis confirmed that the levels of Dp protein were low in lysates from *Dp-/-* fat bodies and from Dp-depleted muscles compared to controls (Figure S3A).

**FIGURE 5:**
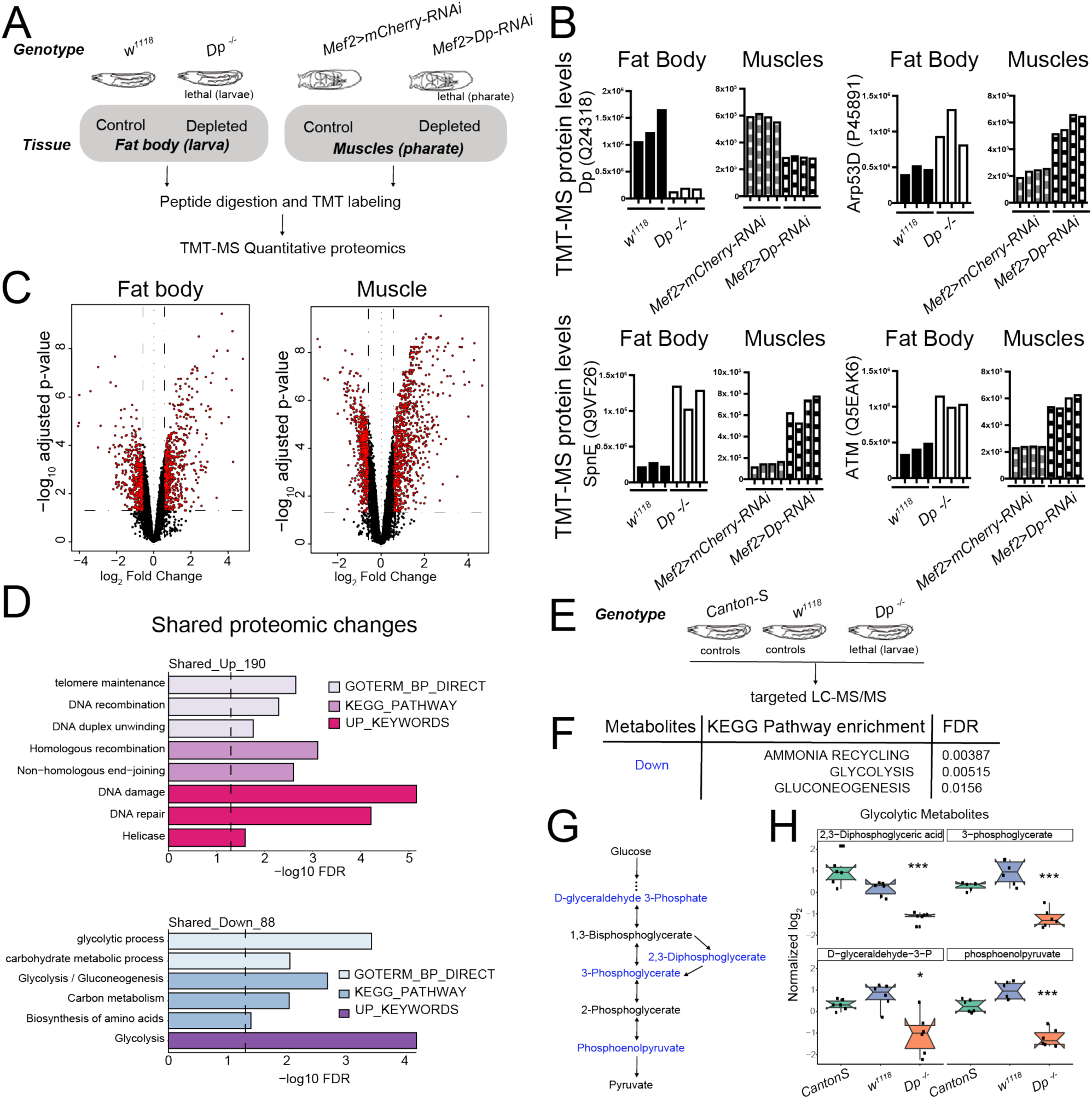
Loss of E2F/Dp impairs carbohydrate metabolism. **A:** Overview of TMT-MS profiles generated from third instar larval fat body in *Dp*-/-mutant and wild type animals and from thoracic muscles in *Mef2>Dp-RNAi* and control animals staged at pharate. 6578 identified proteins in fat bodies and 5730 identified proteins in muscles. **B:** TMT-MS intensities showing protein levels of Dp (Uniprot Q24318), E2F2/DP target protein Arp53 (Uniprot Q9VF26), SpnE (Uniprot P455891) and dATM (Uniprot Q5EAK6) in fat bodies and muscles. Data are represented as individual intensity value for each replicate, n= 3 per genotype in fat body and n=4 per genotype in muscles. **C**: Volcano plots indicating proteins that are differentially expressed between larval WT and *Dp*-/-mutant fat bodies (left panel), and *Mef2>mCherry-RNAi* and *Mef2>Dp-RNAi* pharate muscles (right panel). Significant changes are shown in red (FDR < 0.05 and abs (fold change)>1.5). The x-axis is the log2 of the fold change and the y-axis is the negative log10 of the adjusted p-value. **D:** DAVID functional annotation clustering analysis of proteomic changes in Dp-depleted tissue compared to wild type that are in common between fat body and muscles. Upregulated (top panel, 190 proteins) and downregulated (bottom panel, 88 proteins) were analyzed separately. Dashed line indicates FDR=0.05. Only significant terms (FDR<0.05) are displayed. The categories GO Term biological processes, KEGG pathway and Up keywords are shown. **E**: Overview of targeted LC-MS/MS metabolomic profiles generated from whole third instar larvae. *Dp -/-* mutant animals were compared to two different wild type animals (Canton-S and *w^1118^*). **F:** KEGG pathway enrichment was done on metabolites that were significantly reduced in *Dp -/-* mutant compared to both controls. Only significant terms are shown (FDR<0.05). **G**: Schematic of the flow of glycolysis towards pyruvate. Metabolites that are significantly reduced in *Dp -/-* mutant compared to controls are shown in blue. **H:** Normalized levels of the metabolites D-glyceraldehyde 3-Phosphate, 2,3-Diphosphoglycerate, 3-Phosphoglycerate and Phosphoenolpyruvate in *Dp -/-* mutant compared to controls *w^1118^* and Canton-S. Data are represented as boxplot, which extends from 25 to 75 percentiles; line marks median; whiskers extend to 25/75% plus 1.5 times the interquartile range. Values outside the whiskers are outliers. Welch’s ANOVA test, * p<0.05, *** p<0.001. n=6 per genotype. Full genotypes are (**A**-**D**) *Dp^a2^/Dp^a3^*, *w^1118^*, *UAS-Dp[GD4444]-RNAi,Mef2-GAL4* and *Mef2-GAL4/UAS-mCherry-RNAi* (**E**-**H**) *Dp^Exel7124^/Dp^a3^*, *w^1118^* and *Canton-S*

**SUPPLEMENTARY FIGURE S3:**
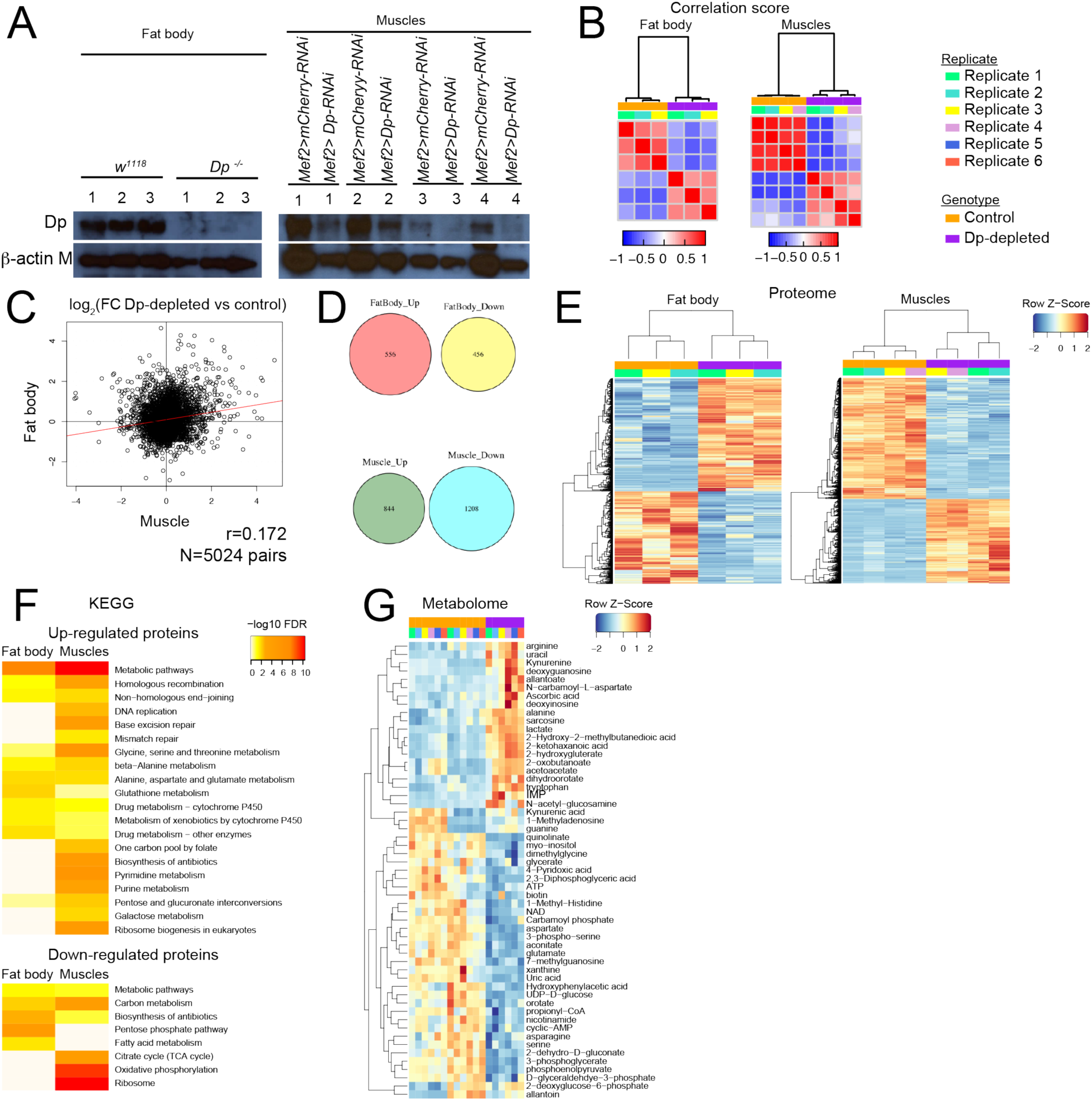
E2F/Dp-deficient muscles and fat bodies undergo severe changes in their proteome and metabolome. **A:** Western blot analysis of fat body (left panel) and muscle (right panel) protein lysates using the antibodies anti-Dp and anti-beta-actin as a loading control. n= 3 replicates per genotype for fat body and n=4 replicates per genotype for muscles. **B**: Correlation score heat map plots of log 2 median protein intensities across replicates from fat body (left panel) and muscle (right panel). **C:** Correlation of Log_2_FC between *Dp*-depleted tissue and wild type in fat body compared to muscles. Pearson correlation test, R=0.172, N=5024 pairs of proteins shared in both tissues. **D**: Venn diagrams: 5024 identified proteins are shared across fat body and muscle proteomes. 556 and 844 proteins are up in *Dp*-depleted compared to control in fat body and muscles, respectively. 456 and 1208 proteins are down in *Dp*-depleted compared to control in fat body and muscles, respectively. **E**: Heatmap representation for hierarchically clustering of differentially expressed proteins (rows) between *Dp*-depleted tissue and control samples (column) in fat body (left panel) and muscle (right panel). n= 3 replicates per genotype for fat body and n=4 replicates per genotype for muscles. Normalized protein levels were clustered in upregulated as FC>=1.5 (red) and downregulated as FC<= −1.5 (blue), moderated *t*-test, FDR<0.05. **F:** Enrichment of KEGG pathway among proteomic changes that are significantly increased (up-regulated, FDR<0.05 and FC>=1.5, top panel) and reduced (downregulated, FDR<0.05 and FC<= −1.5, bottom panel) in *Dp*-depleted tissues compared with control tissue. Fat body is on the left and muscle on the right. **G:** Heatmap representation showing hierarchically clustering of normalized metabolites (row) between *Dp -/-* mutant and both control samples (*Canton-S* and *w^1118^*, column) in third instar larva. Only significant changes to both controls are displayed, ANOVA test, FDR<0.05. n=6 replicates per genotype. Full genotypes are (**A**-**F**) *Dp^a2^/Dp^a3^*,*w^1118^*, *UAS-Dp[GD4444]-RNAi,Mef2-GAL4* and *Mef2-GAL4*/*UAS-mCherry-RNAi* (**G**-**H**) *Dp^Exel7124^*/*Dp^a3^*, *w^1118^* and Canton-S

The changes in the proteomic profiles of fat bodies between the control, *w^1118^*, and *Dp* mutant larva, and in muscles between the control, *Mef2>mCherry*-*RNAi*, and *Mef2>Dp-RNAi* pharates were examined (Sup Table 1). The pairwise correlation between replicate samples were found to be significant (Figure S3B). The mean of correlation values for fat body replicates was r=0.55 for *w^1118^* and r=0.42 for *Dp-/-*, while muscle replicates showed r=0.8 for *Mef2>mCherry*-*RNAi* and r=0.63 for *Mef2>Dp-RNAi*.

In the proteomics datasets, we identified and quantified 6,578 proteins in fat body and 5,370 proteins in muscle samples. We confirmed that the intensity level of Dp protein was downregulated in both *Dp-/-* fat bodies and Dp-depleted muscles compared to the control tissues, respectively. Concordantly, the expression levels of the well-known E2F-direct targets that are repressed by E2F, such as *Arp53D*, *SpnE*, and *dATM*, were upregulated in both *Dp-/*-fat bodies and Dp-depleted muscles (Figure 5B).

A set of 5,024 proteins that were shared between both tissues was selected for integrative analysis (Sup Table 2). A log_2_ fold change (FC) ratio was calculated to identify proteins that were changed upon the loss of *Dp*. These two datasets showed a Pearson correlation of *r* = 0.172 (Figure S3C) indicating that a relevant subset of the changes detected in fat body were also present in muscles. Statistically significant changes were revealed using a false discovery rate (FDR) < 5%, and either a log_2_FC > 0.5 or a log_2_FC < - 0.5 as cut-off values for upregulated and downregulated proteins, respectively (Figure 5C). We found that 556 proteins increased and 456 proteins decreased in *Dp-/-* fat bodies, whereas 844 increased and 1,208 proteins decreased in Dp-depleted muscles (these are visualized in heatmaps in Figure S3D-E).

KEGG pathways enrichment analysis was performed using the functional annotation tool DAVID (Figure S3F, sup table S3) to obtain an overall picture of the changes resulting from the loss of E2F/Dp in the two tissues. Categories related to *DNA repair*, *glutathione metabolism* and *amino acid metabolism* were significantly enriched for upregulated proteins in both tissues, while *nucleotide metabolism* was only significantly enriched in muscle (FDR<5%, Figure S3F, top panel). Similar changes have been linked to the loss of E2F/Dp in previous studies (Guarner et al., 2017; Nicolay et al., 2015, 2013). Additionally, proteins related to *cytochrome p450 enzymes* that catalyze detoxification and biosynthetic reactions in fat body (Chung et al., 2009) were upregulated in *Dp* mutants (Figure S3F, top panel). Among the downregulated proteins, the fat body proteome was enriched for *pentose phosphate pathway* and *fatty acid metabolism*, whereas *citrate cycle*, *oxidative phosphorylation* and *ribosome* categories were significantly overrepresented among the muscle proteome upon the loss of Dp (Figure S3F, bottom panel).

Next, we focused on proteomic changes that were shared between these two tissues since these may reflect an E2F function that is common to both tissues (Figure 5D, sup table S4). As expected, the upregulated proteins in Dp-deficient fat body and muscle showed a significant enrichment for *DNA damage*, *DNA recombination* and *homologous recombination* (Figure 5D, top panel, FDR < 5 %). The top annotation cluster for downregulated proteins displayed a significant enrichment for *glycolysis*, *gluconeogenesis*, and *carbohydrate metabolic process* (Figure 5D, bottom panel, FDR < 5 %), thus indicating that the loss of Dp alters carbohydrate metabolism in both fat body and muscle.

To explore the metabolic defects triggered by the loss of Dp, we used targeted liquid chromatography tandem mass spectrometry (LC-MS/MS) to profile the metabolic changes upon *Dp* loss. Third instar *Dp^-/-^* larva were collected and compared to two wild types strains, *w^1118^* and *Canton S*, to account for differences in the genetic background (Figure 5E, sup table S5). Fifty-five compounds showed significant changes in the *Dp* mutant compared to both controls (FDR < 5 %, Figure S3G, sup table S6). The increased and decreased metabolites were selected and KEGG pathways enrichment analysis was performed. Interestingly, the major metabolic pathways that showed significant enrichment for downregulated compounds were *glycolysis*, *gluconeogenesis* and *ammonia recycling* (FDR < 5 %, Figure 5F, sup table S7), which is largely consistent with the proteome analysis described above. Notably, four metabolites of the core module of the glycolytic pathway (Figure 5G), 2,3-diphosphoglyceric acid, 3-phosphoglycerate, D-glyceraldehyde-3-phosphate and phosphoenolpyruvate, were significantly reduced in *Dp* mutant compared to both controls (Figure 5H).

We conclude that the tissue-specific depletion of *Dp* results in extensive metabolic changes in both fat body and muscle. These changes were evident in proteomic profiles and confirmed by metabolomic profiling. The alterations indicate that E2F-depleted tissues undergo significant changes in carbohydrate metabolism affecting, in particular, glycolytic metabolites.

### Increasing carbohydrates in fly diet rescues the lethality caused by the loss of Dp in fat body

The changes in the proteomic and metabolomic profiles are very interesting but they raised the question of whether the metabolic changes observed contribute to the lethality of *Mef2>Dp*-*RNAi* or *cg>Dp*-*RNAi* animals. Since diet is known to impact metabolic phenotypes, we asked whether varying the levels of carbohydrates, protein and fat in fly food could alter the lethal stage of these animals.

To properly control the food composition and the effect of nutrients we switched to a semi-defined food, made of sucrose, lecithin and yeast, as major sources of carbohydrates, fat and protein, respectively (Lee and Micchelli, 2013). Control diet contained 7.9% carbohydrates, 0.08% fat and 1.9% protein (Sup Table S8). As expected, *Mef2>Dp*-*RNAi* or *cg>Dp*-*RNAi* did not survive on control diet and died at pupal and pharate stage (Figure 6A-B). Nutrient composition was then altered by varying the amount of a single component of the control diet and the viability of *cg>Dp*-*RNAi* and *Mef2>Dp-RNAi* animals relative to the control genotype was scored. Interestingly, while the survival of *Mef2>Dp-RNAi* animals was unaffected by different nutrient composition (Figure 6B, and Sup 4B), *cg>Dp*-*RNAi* were highly sensitive to dietary changes. The increase in protein content had a negative impact on the survival of *cg>Dp-RNAi*, and resulted in significant developmental delay (Figure S4A, left panel). About 83 % of animals showed melanotic masses at larval stages when reared on high protein diet, and only 35 % of third instar larva progressed onto pupa stages and eventually died (Figure S4C, quantified in Figure S4D). The melanotic masses, which were also observed in the *E2f1* mutant larvae (Royzman et al., 1997), are related to the immune system response (Watson et al., 1991). However, reducing the content of protein suppressed the occurrence of melanotic masses (Figure S4C-D), and consequently, increased their survival rate (Figure S4A). In contrast, increasing lipid content up to 1% in diet was beneficial for survival as about 22% of *cg>Dp-RNAi* pupa reached adulthood (Figure S4A, right panel). However, no further rescue in viability was detected beyond 1% lipid. Strikingly, while no *cg>Dp-RNAi* animals survived on control diet (7.9% carbohydrates), almost half of them eclosed when reared in the presence of 12% carbohydrates and the lethality was fully rescued when food contained even higher amount of carbohydrates (16% or 24%) (Figure 6A, left panel). This result was confirmed using a second fat body specific driver *r4-GAL4* (Figure 6A, right panel).

**FIGURE 6:**
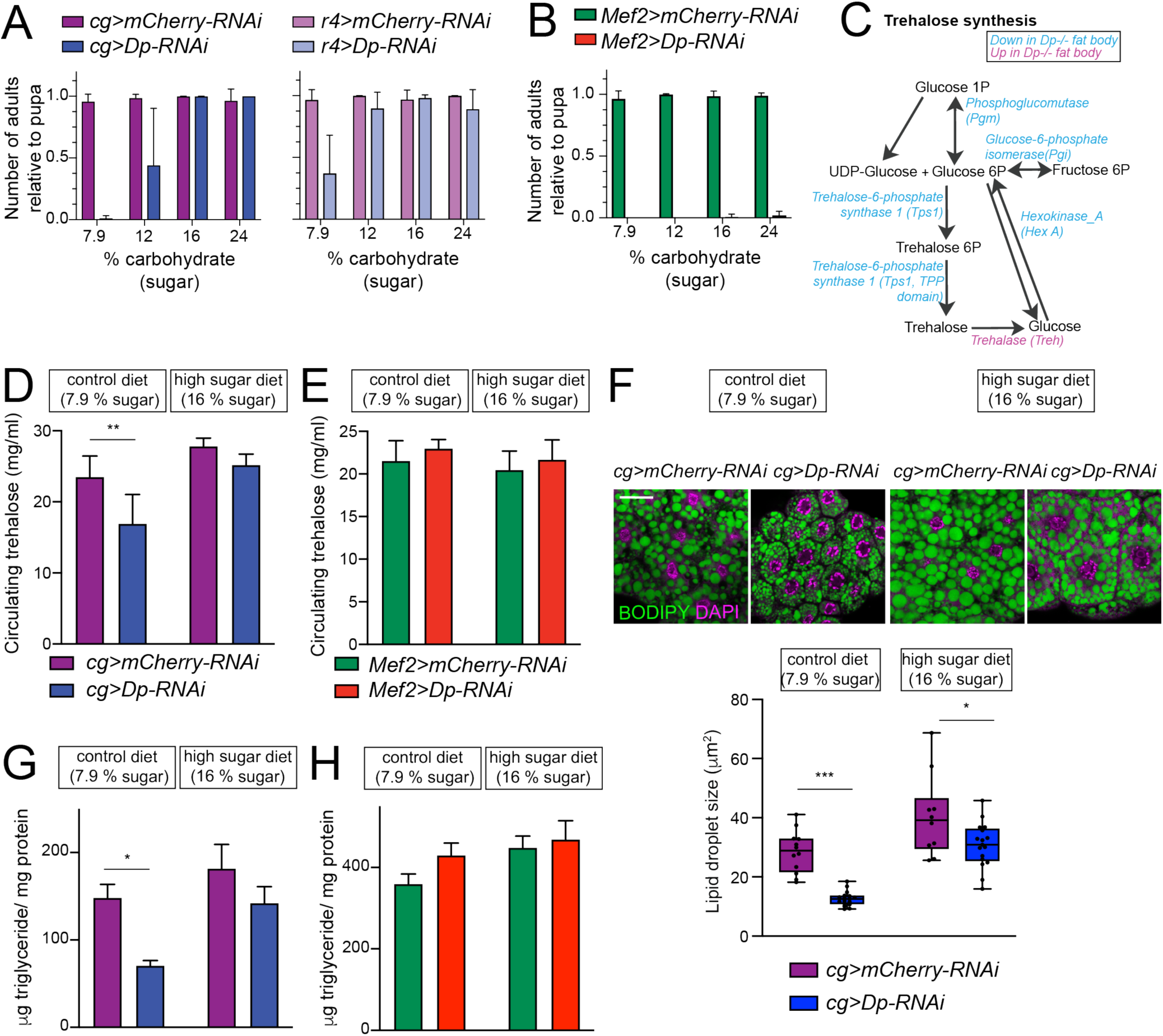
E2F/Dp in fat body exerts systemic effects modulated by sugar supplement. **(A-B)** Number of viable adults (relative to pupa) fed on control diet (7.9% carbohydrate, 0.08% fat and 1.9% protein) and increasing levels of sugar in food (12, 16 and 24% carbohydrate). Data are represented as mean ± SD, n= 6 repeats per condition. (**A**) Left panel: *cg>mCherry RNAi* and *cg>DpRNAi*, right panel: *r4>mCherryRNAi* and *r4>DpRNAi*. (**B**) *Mef2>mCherry-RNAi* and *Mef2>Dp-RNAi* **(C)** Diagram of trehalose synthesis pathway. The enzymes that are significantly downregulated in *Dp*-deficient fat body are indicated in blue, and upregulated in magenta, based on proteome data. **(D-E)** Circulating trehalose levels measured in third instar larval hemolymph. (**D**) *cg>mCherry*-*RNAi* and *cg>Dp-RNAi*, and **(E)** *Mef2>mCherry-RNAi* and *Mef2>Dp-RNAi* larvae fed on control diet (8% carbohydrate) and high sugar diet (16% carbohydrate). Data are represented as mean ± SD, Two-way ANOVA followed by Tukey’s multiple comparisons test, three independent experiments were done, one representative experiment is shown. (**D**) n= 6 per group and ** p = 0.0005. (**E**) n= 3-6 per group and p = 0.5. **(F)** Top panel: Confocal single plane images of third instar larval fat bodies stained with DAPI and BODIPY red. The *cg>Luc-RNAi* and *cg>Dp-RNAi* animals were fed on control diet (7.9% carbohydrate) and supplemented with sugar (16% carbohydrate, high sugar diet). Scale 40 μm. Bottom panel: Measurement of lipid droplet size in fat body. Data are represented as box and whiskers, min to max showing all points, n= 10-17 fat bodies per genotype, Two-way ANOVA followed by Tukey’s multiple comparisons test, * p = 0.02, *** p < 0.0001, three independent experiments, one representative experiment is shown. **(G-H)** Triglycerides content measured in third instar larva and normalized to total protein content. (**G**) *cg>mCherry* RNAi and *cg>DpRNAi*, and **(H)** *Mef2>mCherry RNAi* and *Mef2>DpRNAi* larvae fed on control diet (8% carbohydrate) and high sugar diet (16% carbohydrate). Data are represented as mean ± SEM, two-way ANOVA followed by Tukey’s multiple comparisons test, one representative experiment is shown, (**G**) n= 5-6 per group and * p = 0.004. Three independent experiments were done. (**H**) n= 5-6 per group and p = 0.2, two independent experiments were done. Full genotypes are (**A**) *cg-GAL4,UAS-mCherry-RNAi*, *cg-GAL4/UAS-Dp[GD4444]-RNAi*, *r4-GAL4/UAS-mCherry-RNAi*, *UAS-Dp[GD4444]-RNAi/r4-GAL4* (**B**, **E, H**) *Mef2-GAL4/UAS-mCherry-RNAi*, and *UAS-Dp[GD4444]-RNAi,Mef2-GAL4* (**D, F, G**) cg-*GAL4,UAS-mCherry-RNAi* and *cg-GAL4/UAS-Dp[GD4444]-RNAi*.

**SUPPLEMENTARY FIGURE S4:**
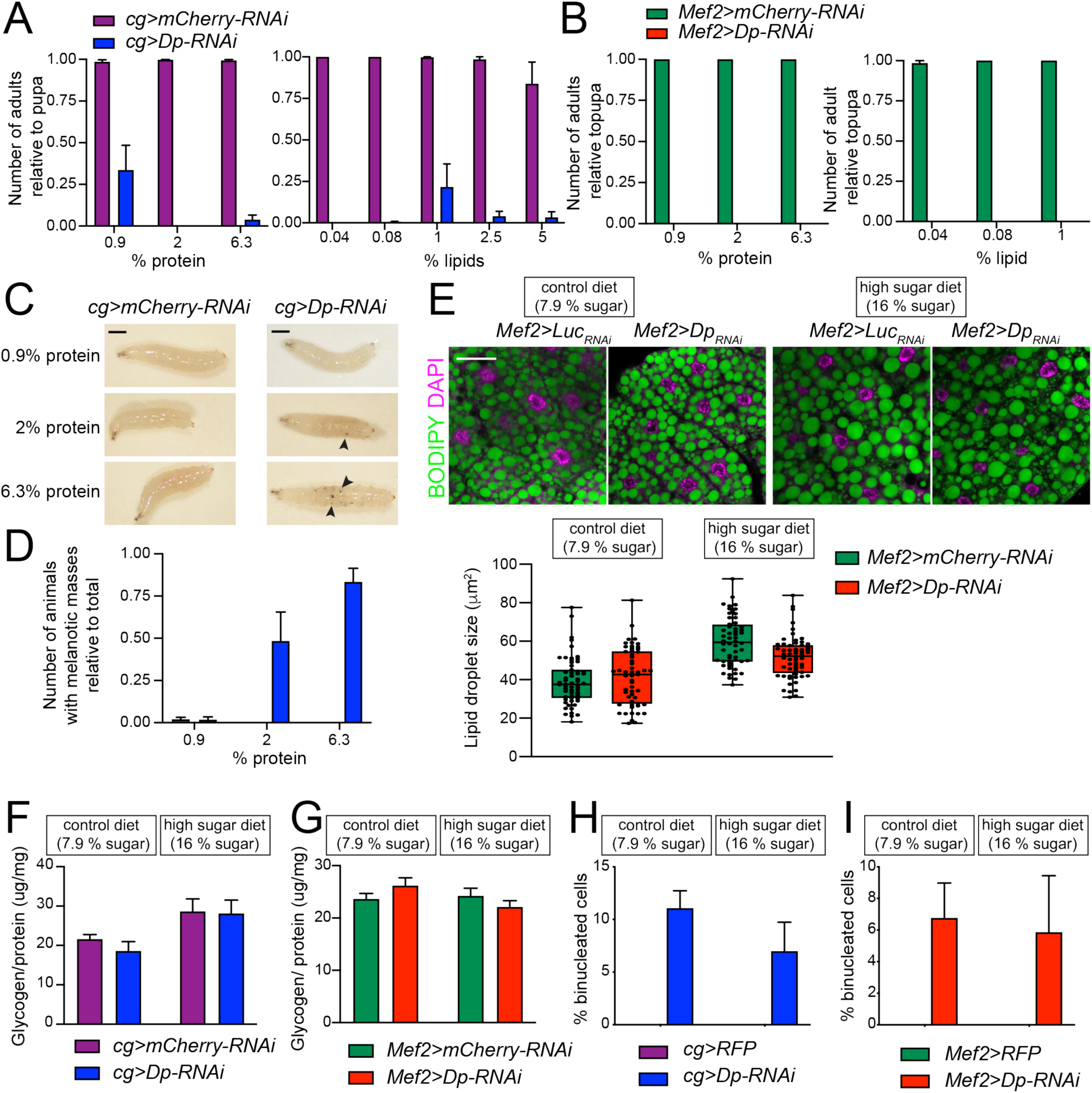
Supplementing food modifies viability of animals with E2F/Dp-deficient fat bodies. **(A-B)** Number of viable adults (relative to pupa) fed on different food conditions. Left panel: increasing levels of protein in food (0.9, 1.96 and 6.3 % protein). Right panel: increasing levels of fat in food (0.04, 0.08, 1.04, 2.5 and 5 % lecithin). Data are represented as mean ± SEM, n= 2-4 sample per condition. Experiment was repeated at least three times. (**A**) *cg>mCherry-RNAi* and *cg>Dp-RNAi*, (**B**) *Mef2>mCherry-RNAi* and *Mef2>Dp-RNAi*. **(C)** Representative images of *cg>mCherry-RNAi* and *cg>Dp-RNAi* larvae fed on increasing levels of protein in food (0.9, 1.96 and 6.3 % protein). Black arrowheads point to melanotic masses. Scale bar 5 mm. **(D)** Quantification of number of larvae fed showing melanotic masses relative to total number of larvae harvested as in C. Data are represented as mean ± SEM, n=2-5. Three independent experiments were done. **(E)** Top panel: Confocal single plane images of third instar larval fat bodies stained with DAPI and BODIPY. The *Mef2>mCherry-RNAi* and *Mef2>Dp-RNAi* animals were fed on control diet (7.9% carbohydrate) and supplemented with sugar (16% carbohydrate, high sugar diet). Scale 40 μm. Bottom panel: Measurement of lipid droplet size in fat body. Data are represented as box and whiskers, min to max showing all points, n= 55-60 images per group in three independent experiments, Two-way ANOVA (main effects only) followed by Tukey’s multiple comparisons test, p= 0.08 for variation between genotypes. **(F-G)** Total whole-body glycogen content normalized to protein content. Third instar larvae **(F)** *cg>mCherry-RNAi* and *cg>Dp-RNAi* (**G**) *Mef2>mCherry-RNAi* and *Mef2>Dp-RNAi* fed on control diet (7.9 % carbohydrate, left panel) and high sugar diet (16% carbohydrate, right panel). Data are represented as mean ± SEM, N= 6 per genotype, three independent experiments were done, Two-way ANOVA followed by Tukey’s multiple comparisons test, (**F**) p = 0.5 and (**G**) p = 0.9 for variation between genotypes. One representative experiment is shown. **(H-I)** Quantification of the percentage of binucleated cells in third instar larval fat bodies. (**H**) *cg>mCherry-RNAi* and *cg>Dp-RNAi* (**I**) *Mef2>mCherry-RNAi* and *Mef2>Dp-RNAi* were fed on control diet (7.9 % carbohydrate) and high sugar diet (16% carbohydrate). Data are represented as mean ± SD, n>200 cells for *Mef2>Dp-RNAi*, and *cg>Dp-RNAi*, and n=200 cells for *Mef2>Luc RNAi* and *cg>Luc RNAi*, which did not show binucleated cells. Full genotypes are (**A, C-D, F**) *cg-GAL4,UAS-mCherry-RNAi*, *cg-GAL4/UAS-Dp[GD4444]-RNAi* (**B, E**) *Mef2-GAL4/UAS-mCherry-RNAi*, *and UAS-Dp[GD4444]-RNAi,Mef2-GAL4* (**H**) *cg-GAL4/UAS-RFP,Dp[GFP], cg-GAL4/UAS-Dp[GFP],UAS-Dp[GD4444]-RNAi*, and (**I**) *UAS-RFP,Dp[GFP];Mef2-GAL4*, and *UAS-Dp[GFP],UAS-Dp[GD4444]-RNAi,Mef2-GAL4*.

One of the functions of fat body is to maintain homeostasis of trehalose, a main circulating sugar in hemolymph (Becker et al., 1996), allowing animals to adapt to a high sugar diet. The synthesis of trehalose occurs in fat body from glucose-6P and is regulated by trehalose-6-phosphate synthase (Tps1) (Figure 6C) (Elbein et al., 2003). We noted that the levels of Tps1 proteins were significantly reduced in *Dp* mutant fat body, along with other enzymes that generate glucose-6P including phosphoglucomutase, glucose-6phosphate isomerase and hexokinase_A in the proteomic datasets (Figure 6C and Table S2). Additionally, Trehalase, an enzyme that converts trehalose back to glucose, was significantly increased in *Dp*-deficient fat body. This suggested that animals lacking Dp in the fat body may be defective in the regulation of trehalose synthesis.

To test this idea directly, trehalose was measured in hemolymph of *cg>Dp-RNAi* and control, *cg>mCherry-RNAi*, third instar larvae fed either a control diet or a high sugar diet containing 16% sugar. Notably, trehalose was significantly reduced in hemolymph of *cg>Dp-RNAi* larva compared to matching control (Figure 6D) when animals were reared on control diet. Switching to high sugar diet significantly improved the levels of trehalose in *cg>Dp-RNAi* to levels comparable to *cg>mCherry-RNAi* larva that were fed a control diet (Figure 6D). In contrast, the levels of circulating trehalose were largely unchanged in *Mef2>Dp-RNAi* compared to control (Figure 6E).

The fat body is the principal site of stored fat in *Drosophila*. *Dp* mutants were previously shown to have lipid droplets of reduced and irregular size (Guarner et al., 2017). Therefore, we examined the impact of high sugar diet on fat body in *cg>Dp-RNAi*. Fat bodies of *cg>Dp-RNAi* and control larva were dissected and stained with BODIPY to visualize lipid droplets. In control diet, Dp-depleted fat bodies displayed smaller lipid droplet size compared to control (Figure 6F, quantification in right panel), which is indicative of defects in fatty acid synthesis. Concordantly, the expression of Acetyl-CoA carboxylase (ACC), the rate-limiting enzyme for fatty acid synthesis, was significantly reduced in fat body of *cg>Dp-RNAi* compared to control (Suppl table S2). As expected (Musselman et al., 2011; Pasco and Léopold, 2012), feeding control larva a high sugar diet resulted in significantly larger lipid storage droplets (Figure 6F). A similar trend was observed in *cg>Dp-RNAi* although the increase in droplet size was not as robust as in the wild type (Figure 6F, quantification in bottom panel). Nevertheless, feeding *cg>Dp-RNAi* animals on high sugar diet dramatically improved the lipid droplet size in comparison to control diet. This was further corroborated by measuring total triglycerides content at third instar larval development (Figure 6G). The levels of triglycerides normalized to total protein content were significantly reduced in *cg>Dp-RNAi* animals reared on control diet compared to *cg>mCherry-RNAi*. Moreover, the normalized levels of triglycerides were fully restored in *cg>Dp-RNAi* animals fed on high sugar diet. Thus, feeding animals with high content of sugar increased their lipid storage, as evidenced by examining both the lipid droplet size in fat body and the total content of triglycerides, and could, subsequently, contribute to rescue animal viability upon *Dp* knockdown in fat body.

In contrast, muscle-specific Dp depletion did not affect the size of lipid droplets in the fat body or triglycerides content of *Mef2>Dp-RNAi* (Figure S4G and Figure 6H). However, the increase of the lipid droplet size in *Mef2>Dp-RNAi* animals reared on high sugar was less than in control fat bodies (Figure S4E, quantification in bottom panel). This suggests that the loss of Dp in muscle impairs the ability of fat body to regulate fat storage.

Dp-depleted fat body failed to maintain the proper level of circulating sugar in hemolymph. Since glucose is converted into glycogen in both fat body and muscle for storage, we measured the level of glycogen in *cg>Dp-RNAi* and in *Mef2>Dp-RNAi*. As shown in Figure S4F-G, there were no major defects in glycogen storage normalized to protein content in *cg>Dp-RNAi* and *Mef2>Dp-RNAi* compared to control animals fed either a control or a high sugar diet.

As mentioned above, Dp depletion in fat body leads to the appearance of binucleated cells (Figure 2A-B) (Guarner et al., 2017). Therefore, we asked whether this phenotype can be suppressed by food composition. Binucleated cells were readily found in both *cg>Dp-RNAi* and *Mef2>Dp-RNAi* larvae fed a control diet, but the phenotype was not modified by rearing larvae on high sugar diet (Figure S4H-I). Since dietary changes suppress the defects and fully rescue animal lethality we infer that the essential function of Dp in fat body is to maintain homeostasis of the circulating sugar trehalose and to regulate fat storage. The fact that binucleated cells persisted in the fat body of rescued animals suggests that the binucleated phenotype is most likely distinct from the metabolic defects.

## DISCUSSION

The experiments described here were prompted by a paradox: if Dp has tissue-specific functions in both the muscle and fat body that are essential for viability, how is it that restoring expression in just one of these tissues can be sufficient to rescue *Dp*-null mutants to viability? We anticipated that *Dp* loss in one of these tissues might cause changes in the other tissue, and/or that the tissue-specific changes might converge through systemic changes. The results described here provide important new insights into the consequences of eliminating E2F/Dp regulation in *Drosophila*. Proteomic and metabolomic profiles reveal extensive changes in *Dp*-depleted tissues and show that the loss of E2F/Dp regulation causes a complex combination of tissue-intrinsic changes and systemic changes. The results described here show that E2F/Dp contributes to animal development and viability by preventing both types of defects.

It is clear from the molecular profiles that *Dp* loss causes extensive metabolic changes in both muscle and fat body. The profiles indicate that E2F/Dp participates, in particular, in the regulation of carbohydrate metabolism in both tissues. Such extensive changes to cellular metabolism likely have biological significance in both contexts, but we know that this is certainly true when *Dp* is depleted in the fat body because the lethality of *cg>Dp-RNAi* could be rescued by increasing dietary sugar. In *Drosophila*, the fat body functions as a key sensor of the nutritional status and it couples systemic growth and metabolism with nutritional availability. The lethality associated with *cg>Dp-RNAi* may, therefore, stem from an imbalance between energy production and systemic growth.

Carbohydrate metabolism defects in *cg>Dp-RNAi* larvae are consistent with studies of mammalian E2F1 in liver and adipose tissue, tissues that metabolize nutrients and stores reserves of lipid and glycogen. E2F1 was shown to be required for the regulation of glycolysis and lipid synthesis in hepatocytes (Denechaud et al., 2016). In adipose tissue E2F1 was implicated in the regulation of PPARγ, the master adipogenic factor, during early stages of adipogenesis (Fajas et al., 2002). Moreover, the gene encoding for the pyruvate dehydrogenase kinase 4 (PDK4), a key nutrient sensor and modulator of glucose oxidation, and 6-phosphofructo-2-kinase/ fructose-2,6-bisphosphatase, a glycolytic enzyme, are directly regulated by E2F1 (Fernandez Mattos et al., 2002; Hsieh et al., 2008). Our results highlight the conserved role of E2F/DP in regulating metabolism. A major function for the fat body is to control whole-animal sugar homeostasis. Interestingly, trehalose and glucose metabolism plays a key role in regulating systemic energy homeostasis (Matsuda et al., 2015; Yasugi et al., 2017). Proper trehalose homeostasis is required for adapting animals to various dietary conditions, as evidenced by *Tps1* mutants, which do not survive in low sugar diet (Matsuda et al., 2015). The fact that the lethality associated with *cg>Dp-RNAi* can be suppressed by a sugar supplement that results in a trehalose increase, underscores the importance of E2F function for animal development, which was not previously appreciated.

In contrast with *cg>Dp-RNAi,* the lethality associated with *Mef2>Dp-RNAi* was not rescued by altering the diet. However, the appearance of binucleated cells in the fat bodies of *Mef2>Dp-RNAi* larvae demonstrates that the muscle-specific knockdown of *Dp* does have systemic effects. These binucleated cells resemble defects triggered by the loss of E2F in the fat body and are also seen in *Dp* mutant larvae. We note that the mechanisms leading to the formation of binucleates in *Mef2>Dp-RNAi* animals may differ from *cg>Dp-RNAi* and *Dp*-/-mutants (Guarner et al., 2017) since they are not associated with a DNA-damage response. Indeed, the binucleated cells in the fat body of *Mef2>Dp-RNAi* animals may be a symptom of stress; evidence for paracrine signals from muscle support this idea (Demontis et al., 2014; Demontis and Perrimon, 2009; Zhao and Karpac, 2017). We infer that the tissue-specific depletion of *Dp* causes systemic effects in both *cg>Dp-RNAi* and *Mef2>Dp-RNAi* larvae, but it is particularly evident in *cg>Dp-RNAi* animals, which have a low level of circulating trehalose in hemolymph.

The level of circulating trehalose is a net result of the amount of trehalose released to hemolymph by fat body and its consumption by other organs, including rapidly growing muscles. The trehalose requirement of wild type organs in *cg>Dp-RNAi* animals may provide an additional challenge to the *Dp*-deficient fat body to maintain trehalose homeostasis. Conversely, the reduced demand for trehalose by *Dp*-deficient organs in the *Dp -/-* mutants may shift the balance towards increased levels of circulating trehalose. Thus, one implication of our work is that the severity of the phenotype may differ between the whole body *Dp-/-* mutant and the tissue-specific *Dp* inactivation.

There is growing evidence of the systemic effects that muscles exert in different settings, including development and aging (Demontis et al., 2014, 2013; Demontis and Perrimon, 2009; Zhao and Karpac, 2017). Myokines are thought to be the main mediators of the inter-tissue communication between muscles and fat body, and other distant tissues. Here, we report that *Dp*-deficient muscle causes the appearance of binucleated cells in the distant fat body of *Mef2>Dp-RNAi* larvae. Our data strongly argue that the fat body phenotype is due to the tissue-specific loss of *Dp* in muscles as the levels of Dp remain unchanged in *Mef2>Dp-RNAi* fat body compared to control animals. This is further validated by lineage tracing, thus confirming the muscle-specific expression of the *Mef2-GAL4* driver. The *Mef2>Dp-RNAi* animals raised on high sugar diet may have a mild defect in the formation of lipid droplets in fat body without an effect on global body triglyceride content. Maintaining proper homeostasis of triglyceride metabolism at an organismal level requires the orchestration of elaborated endocrine mechanism for inter-tissue communication (Heier and Kühnlein, 2018). Myokines and other cytokines, communicate systemic changes in nutrient sensing status and energy substrate storage via cross talk with insulin and Akt signaling pathway (Demontis et al., 2013; Géminard et al., 2009; Zhao and Karpac, 2017).

Since E2F/DP proteins are master regulators of cell proliferation the finding that *Dp* mutant animals develop to the late pupal stages and largely without defects was surprising. It also gave the impression that E2F/DP could be eliminated without major consequences. The results described here paint a different picture. The molecular profiles show extensive changes in *Dp*-deficient tissues. Indeed, 20 % and 51 % of the proteins that we quantified in fat bodies and muscles, respectively, were expressed at statistically different levels following tissue-specific *Dp* knockdown. The second conclusion of our work is that Dp inactivation exerts a systemic effect and may impact distant wild type tissues, which was completely missed in previous analysis of whole body *Dp* mutants and became only evident when *Dp* was inactivated in a tissue-specific manner. We report that *Dp*-deficient muscles affect the morphology of distant fat body, while down regulation of Dp in fat body results in low level of circulating trehalose, which is likely the main cause of lethality of *cg>Dp-RNAi* animals. Thus, the *Dp* mutant phenotype is a complex combination of both tissue intrinsic and systemic effects.

## MATERIALS AND METHODS

### Fly stocks

Flies were raised in vials containing standard cornmeal-agar medium at 25° C. The *w^1118^* flies were used as wildtype (WT) control flies. Either the *Dp^a3^* and *Dp^a4^* alleles or *Dp^a3^* and deficiency *Df(2R)Exel7124*, which deletes the entire Dp gene, were used in this work to obtain the trans-heterozygous *Dp* mutant larvae for proteome and metabolome experiments, respectively (Frolov et al., 2005; Royzman et al., 1997). The trans-heterozygous *E2f2 ^76Q1 /c03344^; E2f1 ^91 / m729^* mutant animals were used as double *E2f1* and *E2f2* mutants. The GAL4 drivers, *P{Cg-GAL4.A}2* and *P{GAL4-Mef2.R}3*, and the following control UAS-RNAi lines from the TRIP collection: *Luciferase* (*P{y[+t7.7] v[+t1.8]=TRiP.JF01355}attP2*) and *mCherry* (*P{VALIUM20-mCherry}attP2*) obtained from Bloomington Drosophila Stock Center (Bloomington, IN, USA). The line *UAS-Dp-RNAi* was obtained from the library RNAi-GD (ID 12722) at the Vienna Drosophila Resource Center (Vienna, Austria). The stock *UAS-G-TRACE* (Evans et al., 2009) was used to trace the expression of the drivers. The *P{PTT-GA}Dp^CA06954^* line from the Carnegie collection (Buszczak et al., 2007), here annotated as DpGFP, contains a GFP-expressing protein trap insertion (Zappia and Frolov, 2016). The *P{UAS-Dp.D}* was used to overexpress *Dp* in the *Dp* mutant background (Du et al., 1996; Neufeld et al., 1998; Zappia and Frolov, 2016).

### Fly viability assay

The total number of pupae, pharate pupae, and adult flies able to eclose out of the pupal case were scored. The pupal developmental stages were assessed by following markers of metamorphosis (Ashburner et al., 2005). At least 58 flies per group were scored in a minimum of three independent experiments.

### Fly food recipes

All flies were raised on Bloomington standard cornmeal food. After eclosion, adults were transferred to different fly food composition, which was made based on the semi-defined control diet (Lee and Micchelli, 2013) with adjustments. Control diet was made of 1 % agar (Lab scientific, Fly 8020), 4.35 % brewers yeast (MP Biomedicals, 2903312), 0.04 % lecithin (soybean, MP Biomedicals, 102147), propionic acid (0.5 %v/v), and 7.9 % sucrose (FCC Food grade, MP Biomedicals, 904713). The adjustments for each food type are detailed in Sup Table S8.

### Hemolymph extraction

Hemolymph was collected from approximately 10 or 15 third instar larvae per sample. Protocol was adapted from (Tennessen et al., 2014). Each animal was rinsed with ddH_2_O, carefully punctured in the mouth hook using a tungsten needle and placed in a 0.5 ml tube with a hole at the bottom of the tube. This tube was then placed in a 1.5 ml tube and centrifuged to max speed for 10 seconds. Approximately 1 µl of hemolymph was collected for each sample. Hemolymph was diluted 1:50 in trehalase buffer (TB) (5 mM Tris pH 7.6, 137 mM NaCl, 2.7 mM KCl). Samples were heat-treated for 5 min at 70 °C and centrifuge for 3 min at max speed at 4°C. Supernatant was quickly snap-frozen and stored at −80 °C until all samples were harvested.

### Trehalose measurement

Hemolymph samples were further diluted to final dilution 1:150 with buffer TB. Circulating trehalose was measured in hemolymph as previously described (Tennessen et al., 2014). Briefly, an aliquot of each sample was treated with porcine trehalase (Sigma, T8778-1UN) overnight at 37 °C in a G1000^TM^ Thermal cycler (BIO-RAD) to digest trehalose to produce free glucose. In parallel, another aliquot was incubated with buffer TB to determine the levels of glucose. The total amount of glucose was determined using the glucose (HK) assay reagent (Sigma, G3293) following a 15 min incubation at room temperature. Trehalose (Sigma, 90208) and glucose standard solutions (Sigma, G3285) were used as standards. Plate reader BioTek Epoch was used to read absorbance at 340mm. The trehalose concentrations for each sample were determined by subtracting the values of free glucose in the untreated samples. Each sample was measured twice, a total of 6 independent biological samples were collected by group and three independent experiments was done.

### Quantification of glycogen and protein content

Third instar larvae were harvested to measure glycogen as previously described (Tennessen et al., 2014). Briefly, seven animals were collected per sample, rinsed with ddH_2_O and homogenized in 100 µl PBS 1x. Samples were heat-treated at 70 °C for 10 minutes and centrifuged at maximum speed for 3 minutes at 4 °C. Supernatant was stored at −80 °C until all samples were collected. Samples were diluted 1:6 in PBS 1x for the assay and transferred to two wells. One well was treated with amyloglucosidase (Sigma A1602) and the second well with PBS 1x. The plate was incubated at 37 °C for 1 hour. Then, the total amount of glucose was determined using 100 µl of glucose (HK) assay reagent (Sigma, G3293) following a 15 min incubation at room temperature. Glycogen and glucose standard solutions were used as standards. Plate reader BioTek Epoch was used to read absorbance at 340mm. The glycogen concentrations for each sample were determined by subtracting the values of free glucose in the untreated samples. Each sample was measured twice, a total of 6 independent biological samples were collected by group and three independent experiments was done. Total glycogen was normalized to soluble protein amount. Aliquots of larval homogenate were removed prior heat-treatment to measure soluble protein using a Bradford assay (Bio-Rad 500-0006) with BSA standard curves.

### Triglycerides quantification

A coupled colorimetric assay was used to quantify triglycerides by measuring free glycerol as previously described (Tennessen et al., 2014). Briefly, seven animals were collected per sample, rinsed with ddH_2_O and homogenized in cold 100 µl PBS-T (PBS 0.05% Tween-20). Samples were heat-treated at 70 °C for 10 minutes and stored at −80 °C until all samples were collected. Samples were diluted 1:6 in PBS-T for the assay and transferred to two wells. One well was treated with Triglyceride reagent (Sigma, T2449) and the second well with PBS-T. The plate was incubated at 37 °C for 30 min in a G1000^TM^ Thermal cycler (BIO-RAD). Then, the total amount of free glycerol was determined using the 100 µl free glycerol reagent (Sigma, F6428) following a 5 min incubation at 37 °C. Glycerol (triolein, Sigma, G7793) standard solution was used as standard. Plate reader BioTek Epoch was used to read absorbance at 540mm. The triglyceride concentrations for each sample were determined by subtracting the values of free glycerol in the untreated samples. Each sample was measured twice, a total of 6 independent biological samples were collected by group and three independent experiments was done. Total triglyceride was normalized to soluble protein amount as described above.

### Immunofluorescence

Tissues were dissected and fixed in 4 % formaldehyde in PBS for 30 min. Then, tissues were permeabilized during 10 or 15 minutes in 0.1 % Triton X-100 in PBS or in 0.3 % Triton X-100 in PBS for muscles tissues. Tissues were washed and blocked in 1 % or 2 % BSA PBS. Primary antibodies were incubated overnight at 4 °C in 2 % BSA and 0.1 % Triton X-100 in PBS. After washing three or four times for 10 min each in 0.1 % Triton X-100 (in PBS), secondary antibodies (Alexa Fluor, Cy3-or Cy5-conjugated anti-mouse and anti-rabbit secondary antibodies, Life Technologies and Jackson Immunoresearch Laboratories) were incubated for 60 or 90 min in 10 % normal goat serum 0.1 % Triton X-100 in PBS. After washing three times with 0.1 % Triton X-100 (in PBS), tissues were mounted on glass slides in glycerol with antifade or in Vectashield with DAPI (Vector Laboratories). All steps were performed at room temperature, unless otherwise stated.

In the case of fat bodies the fixation was done for 60 min and PBS 1x was used throughout the protocol instead of 0.1 % Triton X-100 in PBS 1x (Guarner et al., 2017). For transverse plane sectioning of thoraces staged at 96 h flies were snap-frozen in liquid nitrogen, cut twice with a razor, and fixed for 1 h in relaxing buffer (Zappia and Frolov, 2016). For larval body wall musculature staining, larva was dorsally opened, pinned in a Sylgard dish and fixed for 20 min. A minimum of five to eight animals per genotype was dissected per experiment, and the staining was carried out two or three times.

The primary antibodies were mouse anti-ß PS-integrin (CF.6G11, 1:50, DSHB), mouse monoclonal anti-Dp antibody (Yun6, dil 1:10,(Du et al., 1996)) used in fat bodies and rabbit polyclonal anti-Dp antibodies (#212, dil 1:300 (Dimova et al., 2003)) used in muscles, anti-GFP (FITC, 1:500, Abcam ab6662), Guinea pig anti-Rad50 (1:100, (Gao et al., 2009)), Guinea pig anti-Mre11 (1:100, (Gao et al., 2009)). Rhodamine–phalloidin or fluorescein isothiocyanate–phalloidin were used to counterstain, and 4,6-diamidino-2-phenylindole (DAPI) for nucleus staining.

### Lipid droplet detection

To visualize lipid droplets, dissected third instar larval fat bodies were fixed in 4 % formaldehyde in PBS for 1h at room temperature and washed three times in PBS 1X. Fat bodies were incubated in solution conatining both 0.5 µg/ml BODIPY 493/503 (Invitrogen, D3922) and DAPI diluted in PBS 1x, for 10 minutes at room temperature, then washed three times in PBS 1X.

### Confocal microscopy/ Image acquisition

Fluorescent images were acquired with the laser-scanning confocal microscope (Zeiss LSM 700) using x 20/0.8, and x 40/1.2 objectives at University of Illinois at Chicago and 710 Zeiss Confocal microscope at MGH Cancer Center. Images were processed using Image J (1.52k5, National Institutes of health, USA) and Photoshop CC 2019 (Adobe Systems). All images are confocal single-plane images. Only representative images are shown.

### Quantitative Proteomics

#### Sample preparation

Fat bodies and thoracic muscles were dissected in cold PBS 1x. To pellet the dissected tissues, vials were centrifuged at 4 °C at max speed, and PBS was removed prior to snap-freezing. Collected tissues were thaw and resuspended in modified protein lysis buffer (50 mM HEPES pH 8, 100 mM KCl, 2 mM EDTA, 10 mM NaF, 10% Glycerol, 0.1% NP-40, 1 mM dithiothreitol (DTT), 1 mM PMSF and Roche protease inhibitors) and homogenized on ice. The amount of total protein was measured with Lowry colorimetric assay (DC, Bio-Rad) for fat bodies and Bradford standard assay (Bio-Rad 500-0006) for muscles. Western blotting was carried out using standard procedures. The mouse anti-DP Yun (#6, 1:5 (Du et al., 1996) and the mouse beta actin (1:1000, Abcam, Cat#ab8224) antibody was used as loading control in western blot assays.

#### Multiplexed quantitative mass spectrometry-based proteome

The TMT-10 plex reagents and the SPS-MS3 method on an Orbitrap Fusion mass spectrometer (Thermo Scientific) (Edwards and Haas, 2016; Guarner et al., 2017; Lapek et al., 2017; McAlister et al., 2014) were used to profile *w^1118^* and *Dp -/-* whole larval lysates, and *Mef2>mCherry-RNAi* and *Mef2>Dp-RNAi* thoracic muscle lysates in triplicate and quadruplet, respectively. Disulfide bonds were reduced, free thiols were alkylated with iodoacetamide; proteins were purified by MeOH/CHCl3 precipitation and digested with Lys-C and trypsin, and peptides were labeled with TMT-10plex reagents (Thermo Scientific) (Edwards and Haas, 2016; McAlister et al., 2014). Labeled peptide mixtures were pooled and fractionated by basic reversed-phase HPLC. Four fractions were analyzed by multiplexed quantitative proteomics performed on an Orbitrap Fusion mass spectrometer (Thermo Scientifc) using a Simultaneous Precursor Selection (SPS) based MS3 method (McAlister et al., 2014). MS2 spectra were assigned using a SEQUEST-based proteomics analysis platform (Huttlin et al., 2010). The protein sequence database for matching the MS2 spectra was based on v5.57 of the *D. melanogaster* proteome retrieved from Flybase (Attrill et al., 2016). Peptide and protein assignments were filtered to a false discovery rate of < 1 % employing the target-decoy database search strategy (Elias and Gygi, 2007) and using linear discriminant analysis and posterior error histogram sorting (Huttlin et al., 2010). Peptides with sequences contained in more than one protein sequence from the UniProt database were assigned to the protein with most matching peptides (Huttlin et al., 2010). We extracted TMT reporter ion intensities as those of the most intense ions within a 0.03 Th window around the predicted reporter ion intensities in the collected MS3 spectra. Only MS3 with an average signal-to-noise value of larger than 20 per reporter ion as well as with an isolation specificity (Ting et al., 2011) of larger than 0.75 were considered for quantification. A two-step normalization of the protein TMT-intensities was performed by first normalizing the protein intensities over all acquired TMT channels for each protein based on the median average protein intensity calculated for all proteins. To correct for slight mixing errors of the peptide mixture from each sample a median of the normalized intensities was calculated from all protein intensities in each TMT channel and the protein intensities were normalized to the median value of these median intensities.

#### Proteomics analysis

A total of 6578 and 5730 proteins in fat body and muscles, respectively, were quantified across all experimental conditions. Only shared proteins between fat bodies and muscles were selected for an integrative analysis, all downstream analysis were done on combined 5024 proteins. Differential protein expressions between *w^1118^* and *Dp-/-* fat bodies and between *Mef2>mCherry-RNAi* and *Mef2>Dp-RNAi* proteomes, were calculated using a moderated T-test. The Benjamini-Hochberg multiple hypothesis correction was applied to calculate corrected p-values (FDR). Differential expression of proteins was considered significant with an FDR < 5 % and an absolute fold change greater than 1.5. <u>Functional Enrichment Analysis</u>: Functional Annotation Clustering of the differentially expressed proteins were analyzed using DAVID platform (https://david.ncifcrf.gov/summary.jsp (Huang et al., 2009a, 2009b)). Functional terms related to biological process (BP), KEGG, and UP_Keyword were identified using FDR <0.05 in the selected top cluster. Furthermore, gene ontology enrichment was analyzed by selecting KEGG pathways and using FDR < 5 %.

### Metabolomics profiles

#### Pre-extraction of metabolites from whole larvae and 13C labeling of whole larvae

Six biological samples per genotype were processed exactly as previously published (Nicolay et al., 2013). From each of the vials that contained ∼30 larvae, animals were isolated and washed twice in ddH2O to remove any excess foodstuff, outside unlabeled metabolites, or excess 13C labeled glutamine. Then animals were collected in 1.5mL tubes and the total weight of each collection of starting material was ∼10 mg to achieve detection of unstable metabolites. ∼3mg of starting material was sufficient for most metabolites. For each condition tested, metabolites were extracted from 6-8 biological replicates of pooled animals from each genotype. Samples were then snap frozen in liquid nitrogen and either stored at −80°C for further processing or processed immediately.

Snap-frozen samples were kept on dry ice during extraction. 500 µL of −80°C MeOH:H_2_O (80:20) was added to each pellet. Pellets were homogenized by hand with a pestle using 3-5 strokes. Samples were 7 vortexed at 4 °C for 1 min and left at −80 °C for 4 hrs. After 4 hrs, samples were vortexed at 4°C for 30 seconds. Samples were clarified at 20K x g, for 0.25hr at 4 °C. Clarified supernatant was transferred to a new 1.5mL tube and stored at −80 °C. Each pellet was reextracted with −80 °C MeOH:H2O (80:20), vortexed for 30 seconds at 4 °C and stored at −80 °C for 0.5hr. Reextracted material was vortexed for 30 seconds at 4 °C and then clarified at 20K x g, for 0.25hr at 4 °C. Clarified supernatants were combined and clarified one more time. Combined supernatants were then evaporated by SpeedVac, snap frozen in liquid nitrogen, and stored at −80 °C. Prior to mass spectrometry analysis, samples were resuspended using 20 µL HPLC grade water.

#### LC-MS/MS

LCMS methodology was performed as described in (Nicolay et al., 2015). In brief, nanospray HPLC-MS was carried out with an Agilent 1260 Infinity pump coupled to a FAMOS+ autosampler and an Exactive Orbitrap mass spectrometer. The mass spectrometer was equipped with an electrospray ionization source operated in negative mode. The mass spectrometer was calibrated using a negative ion calibration solution (Pierce 88324) and the optimized conditions were spray voltage 1.8 kV, spray current 2.1 uA, capillary temperature 301 °C, capillary voltage −52.5, tube lens voltage −150, skimmer voltage −42. The mass spectrometer was run in full scan mode (80-1000 m/z range) with an R= 100,000 at 1 Hz (1 scan/second) with the use of the ion pairing reagent, Tributylamine (Sigma 471313). The stationary phase was a C18 medium (3 µm, 200A) from Maccel. The LC method used was as follows: 0 min, 0% B; 11 min, 5% B; 24 min, 100% B; 30 min, 100% B; 31 min, 0% B, 40 min, 0% B. Injection volume was 1 µL. Flow rate at column bed was 400 nL/min. Buffer A: 5% Methanol, 10 mM TBA, 10 mM Acetic Acid. Buffer B: Methanol. Raw data files were transformed and analyzed in MAVEN (Clasquin et al., 2012; Melamud et al., 2010).

#### Metabolomics analysis

For analyses of metabolite pools the free portal Metaboanalyst (www.metaboanalyst.ca) was used. Raw metabolite measurement data were converted to achieve a normal distribution of the data. For each metabolite, data were median-centered, then log2-transformed across the genotypes followed by autoscaling. Among the 258 metabolites only 210 had at least three strong peaks (out of the 6-8 per group) in each of the genotypes. Analysis was done using data from the 210 metabolites that gave reproducible peaks above noise. Standard compound names were used and the compound pathway library was *Drosophila melanogaster*. Following normalization, altered metabolite levels were functionally compared across all three genotypes. Only metabolites that were significantly altered in the *Dp-/-* mutant when compared to both the Canton S and *w^1118^* control genotypes were considered in the analysis to account for the variation on genetic background. KEGG enrichment analysis was done using metabolites changed up and down in *Dp -/-* mutant to determine what metabolic pathways were altered. Direct comparisons between normalized values of specific metabolites were done using Excel and significance was tested using one-way ANOVA analysis followed by Tukey’s post-hoc analysis.

### Quantification and statistical analysis

#### Image analysis

Muscle area in DLM 4 cross-section of thoracic muscles and body wall muscle VL3 were measured for each animal using ImageJ.

The size of lipid droplets were quantified using the “analyze particles” function of ImageJ. The number of binucleated cells in fat bodies were manually scored on the microscope and normalized to the total number of nuclei counted per field.

The ratio of Dp (and DpGFP) signal relative to nuclear area was calculated using Fiji (https://fiji.sc/) in fat bodies and ImageJ (https://imagej.nih.gov/ij/) in muscles. Measurements for both parameters (raw intensity and nuclear area) were collected simultaneously in individual images.

Scripts used in ImageJ for automatic and unbiased quantification of lipid droplets is included in Source Data. All statistics and graphs were generated with the GraphPad Prism version 9.0.1 (Graphpad Software). The group means were analyzed for overall statistical significance using non-parametric test, including Kruskal-Wallis test followed by Dunn’s multiple comparisons test and Mann-Whitney test, and Two-way ANOVA followed by Tukey’s multiple comparisons test in control and high sugar diet experiments. Both a Spearman’s test for heteroscedasticity and a Kolmogorov-Smirnov and a Shapiro-Wilk (W) test for normality were assessed before choosing Two-way ANOVA statistical analysis. Details on the sample size, number of independent experiment and statistical analysis are listed in Figure legends.

#### Proteomics and metabolomics analysis

All plots for proteomic and metabolomic were generated using R. Further details on the analysis can be found in the proteome and metabolome sections in materials and methods.

### Data availability

All mass spectrometer RAW files for quantitative proteomics analysis can be accessed through the MassIVE data repository (massive.ucsd.edu) under the accession number MSV000086854.

## Acknowledgements

We thank Kristin White and Jason Tennessen for helpful discussions, Isabel Liseth for technical help. We are grateful to the Bloomington Drosophila Stock Center (supported by NIH grant P40OD018537), the Vienna Drosophila Resource Center, the TRiP at Harvard Medical School for fly stocks; the Developmental Studies Hybridoma Bank (DSHB) for antibodies; and to Flybase for online resources on the Database of Drosophila Genes & Genomes. This work was supported by NIH grant R35GM131707 (to M.V.F.) and R01GM117413 (to N.J.D.).

## Author contributions

Conceptualization, M.P.Z., A.G., N.J.D. and M.V.F.; Investigation, M.P.Z., A.G., N.K-S., A.R., B.N., and M.B.; Methodology and Formal Analysis, M.P.Z., A.G., R.M., M.B., W.H., B.N.; Funding Acquisition, N.J.D., W.H., and M.V.F.; Writing – Original Draft,, M.P.Z. and A.G.; Writing –Review & Editing, M.P.Z., N.J.D. and M.V.F; Supervision, W.H., N.J.D. and M.V.F.

## REFERENCE

Annicotte, J.S., Blanchet, E., Chavey, C., Iankova, I., Costes, S., Assou, S., Teyssier, J., Dalle, S., Sardet, C., Fajas, L., 2009. The CDK4-pRB-E2F1 pathway controls insulin secretion. Nat. Cell Biol. 11, 1017–1023. https://doi.org/10.1038/ncb1915

Ashburner, M., Golic, K.G., Hawley, R.S., 2005. Drosophila: A Laboratory Handbook, Second Edi. ed. Cold Spring Harbor Laboratory Press, Cold Spring Harbor, New York.

Attrill, H., Falls, K., Goodman, J.L., Millburn, G.H., Antonazzo, G., Rey, A.J., Marygold, S.J., 2016. Flybase: Establishing a gene group resource for Drosophila melanogaster. Nucleic Acids Res. 44, D786–D792. https://doi.org/10.1093/nar/gkv1046

Becker, A., Schlöder, P., Steele, J.E., Wegener, G., 1996. The regulation of trehalose metabolism in insects. Experientia 52, 433–439. https://doi.org/10.1007/BF01919312

Blanchet, E., Annicotte, J.-S., Lagarrigue, S., Aguilar, V., Clapé, C., Chavey, C., Fritz, V., Casas, F., Apparailly, F., Auwerx, J., Fajas, L., 2011. E2F transcription factor-1 regulates oxidative metabolism. Nat. Cell Biol. 13, 1146–52. https://doi.org/10.1038/ncb2309

Buszczak, M., Paterno, S., Lighthouse, D., Bachman, J., Planck, J., Owen, S., Skora, A.D., Nystul, T.G., Ohlstein, B., Allen, A., Wilhelm, J.E., Murphy, T.D., Levis, R.W., Matunis, E., Srivali, N., Hoskins, R.A., Spradling, A.C., 2007. The carnegie protein trap library: a versatile tool for Drosophila developmental studies. Genetics 175, 1505–31. https://doi.org/10.1534/genetics.106.065961

Chung, H., Sztal, T., Pasricha, S., Sridhar, M., Batterham, P., Daborn, P.J., 2009. Characterization of Drosophila melanogaster cytochrome P450 genes. Proc. Natl. Acad. Sci. U. S. A. 106, 5731–5736. https://doi.org/10.1073/pnas.0812141106

Clasquin, M.F., Melamud, E., Rabinowitz, J.D., 2012. LC-MS Data Processing with MAVEN: A Metabolomic Analysis and Visualization Engine, in: Curr Protoc Bioinformatics. pp. 1–7. https://doi.org/10.1002/0471250953.bi1411s37.LC-MS

Demontis, F., Patel, V.K., Swindell, W.R., Perrimon, N., 2014. Intertissue control of the nucleolus via a myokine-dependent longevity pathway. Cell Rep. 7, 1481–94. https://doi.org/10.1016/j.celrep.2014.05.001

Demontis, F., Perrimon, N., 2009. Integration of Insulin receptor/Foxo signaling and dMyc activity during muscle growth regulates body size in Drosophila. Development 136, 983– 93. https://doi.org/10.1242/dev.027466

Demontis, F., Piccirillo, R., Goldberg, A.L., Perrimon, N., 2013. The influence of skeletal muscle on systemic aging and lifespan. Aging Cell 12, 943–949. https://doi.org/10.1111/acel.12126

Denechaud, P.-D., Fajas, L., Giralt, A., 2017. E2F1, a Novel Regulator of Metabolism. Front. Endocrinol. (Lausanne). 8. https://doi.org/10.3389/fendo.2017.00311

Denechaud, P.D., Lopez-Mejia, I.C., Giralt, A., Lai, Q., Blanchet, E., Delacuisine, B., Nicolay, B.N., Dyson, N.J., Bonner, C., Pattou, F., Annicotte, J.S., Fajas, L., 2016. E2F1 mediates sustained lipogenesis and contributes to hepatic steatosis. J. Clin. Invest. 126, 137–150. https://doi.org/10.1172/JCI81542

Dimova, D.K., Stevaux, O., Frolov, M. V, Dyson, N.J., 2003. Cell cycle-dependent and cell cycle-independent control of transcription by the Drosophila E2F/RB pathway. Genes Dev. 17, 2308–20. https://doi.org/10.1101/gad.1116703

Du, W., Xie, J.-E., Dyson, N.J., 1996. Ectopic expression of dE2F and dDP induces cell proliferation and death in the Drosophila eye. EMBO J. 15, 3684–3692.

Dyson, N.J., 2016. RB1: A prototype tumor suppressor and an enigma. Genes Dev. 30, 1492– 1502. https://doi.org/10.1101/gad.282145.116

Edwards, A., Haas, W., 2016. Multiplexed quantitative proteomics for high-throughput comprehensive proteome comparisons of human cell lines. Methods Mol. Biol. 1394, 1– 13. https://doi.org/10.1007/978-1-4939-3341-9_1

Elbein, A.D., Pan, Y.T., Pastuszak, I., Carroll, D., 2003. New insights on trehalose: A multifunctional molecule. Glycobiology 13, 17–27. https://doi.org/10.1093/glycob/cwg047

Elias, J.E., Gygi, S.P., 2007. Target-decoy search strategy for increased confidence in large-scale protein identifications by mass spectrometry. Nat. Methods 4, 207–214. https://doi.org/10.1038/nmeth1019

Evans, C.J., Olson, J.M., Ngo, K.T., Kim, E., Lee, N.E., Kuoy, E., Patananan, A.N., Sitz, D., Tran, P.T., Do, M.T., Yackle, K., Cespedes, A., Hartenstein, V., Call, G.B., Banerjee, U., 2009. G-TRACE: Rapid Gal4-based cell lineage analysis in Drosophila. Nat. Methods 6, 603–605. https://doi.org/10.1038/nmeth.1356

Fajas, L., Landsberg, R.L., Huss-Garcia, Y., Sardet, C., Lees, J.A., Auwerx, J., 2002. E2Fs regulate adipocyte differentiation. Dev. Cell 3, 39–49. https://doi.org/10.1016/S1534-5807(02)00190-9

Fernandez Mattos, S., Lam, E.W.F., Tauler, A., 2002. An E2F-binding site mediates the activation of the proliferative isoform of 6-phosphofructo-2-kinase/fructose-2,6-bisphosphatase by phosphatidylinositol 3-kinase. Biochem. J. 368, 283–291.

Frolov, M. V, Huen, D.S., Stevaux, O., Dimova, D., Balczarek-Strang, K., Elsdon, M., Dyson, N.J., 2001. Functional antagonism between E2F family members. Genes Dev. 15, 2146–2160. https://doi.org/10.1101/gad.903901

Frolov, M. V, Moon, N.-S., Dyson, N.J., 2005. dDP Is Needed for Normal Cell Proliferation. Mol. Cell. Biol. 25, 3027–3039. https://doi.org/10.1128/MCB.25.8.3027

Gao, G., Bi, X., Chen, J., Srikanta, D., Rong, Y.S., 2009. Mre11-Rad50-Nbs complex is required to cap telomeres during Drosophila embryogenesis. Proc. Natl. Acad. Sci. U. S. A. 106, 10728–10733. https://doi.org/10.1073/pnas.0902707106

Géminard, C., Rulifson, E.J., Léopold, P., 2009. Remote Control of Insulin Secretion by Fat Cells in Drosophila. Cell Metab. 10, 199–207. https://doi.org/10.1016/j.cmet.2009.08.002

Guarner, A., Morris, R., Korenjak, M., Boukhali, M., Zappia, M.P., Van Rechem, C., Whetstine, J.R., Ramaswamy, S., Zou, L., Frolov, M. V., Haas, W., Dyson, N.J., 2017. E2F/DP Prevents Cell-Cycle Progression in Endocycling Fat Body Cells by Suppressing dATM Expression. Dev. Cell 0, 689–703. https://doi.org/10.1016/j.devcel.2017.11.008

Heier, C., Kühnlein, R.P., 2018. Triacylglycerol Metabolism in Drosophila melanogaster 210, 1163–1184. https://doi.org/10.1534/genetics.118.301583

Hsieh, M.C.F., Das, D., Sambandam, N., Zhang, M.Q., Nahle, Z., 2008. Regulation of the PDK4 Isozyme by the Rb-E2F1 Complex. J. Biol. Chem. 283, 27410–27417. https://doi.org/10.1074/jbc.M802418200

Huang, D.W., Lempicki, R. a, Sherman, B.T., 2009a. Systematic and integrative analysis of large gene lists using DAVID bioinformatics resources. Nat. Protoc. 4, 44–57. https://doi.org/10.1038/nprot.2008.211

Huang, D.W., Sherman, B.T., Lempicki, R.A., 2009b. Bioinformatics enrichment tools: Paths toward the comprehensive functional analysis of large gene lists. Nucleic Acids Res. 37, 1–13. https://doi.org/10.1093/nar/gkn923

Huttlin, E.L., Jedrychowski, M.P., Elias, J.E., Goswami, T., Rad, R., Beausoleil, S.A., Villén, J., Haas, W., Sowa, M.E., Gygi, S.P., 2010. A tissue-specific atlas of mouse protein phosphorylation and expression. Cell 143, 1174–1189. https://doi.org/10.1016/j.cell.2010.12.001

Lapek, J.D., Greninger, P., Morris, R., Amzallag, A., Pruteanu-Malinici, I., Benes, C.H., Haas, W., 2017. Detection of dysregulated protein-association networks by high-throughput proteomics predicts cancer vulnerabilities. Nat. Biotechnol. 35, 983–989. https://doi.org/10.1038/nbt.3955

Lee, W.C., Micchelli, C.A., 2013. Development and Characterization of a Chemically Defined Food for Drosophila. PLoS One 8, 1–10. https://doi.org/10.1371/journal.pone.0067308

Matsuda, H., Yamada, T., Yoshida, M., Nishimura, T., 2015. Flies without trehalose. J. Biol. Chem. 290, 1244–1255. https://doi.org/10.1074/jbc.M114.619411

McAlister, G.C., Nusinow, D.P., Jedrychowski, M.P., Wühr, M., Huttlin, E.L., Erickson, B.K., Rad, R., Haas, W., Gygi, S.P., 2014. MultiNotch MS3 enables accurate, sensitive, and multiplexed detection of differential expression across cancer cell line proteomes. Anal. Chem. 86, 7150–7158. https://doi.org/10.1021/ac502040v

Melamud, E., Vastag, L., Rabinowitz, J.D., 2010. Metabolomic analysis and visualization engine for LC - MS data. Anal. Chem. 82, 9818–9826. https://doi.org/10.1021/ac1021166

Musselman, L.P., Fink, J.L., Narzinski, K., Ramachandran, P.V., Hathiramani, S.S., Cagan, R.L., Baranski, T.J., 2011. A high-sugar diet produces obesity and insulin resistance in wild-type Drosophila. Dis. Model. Mech. 4, 842–849. https://doi.org/10.1242/dmm.007948

Neufeld, T.P., de la Cruz, A.F.A., Johnston, L.A., Edgar, B.A., 1998. Coordination of growth and cell division in the Drosophila wing. Cell 93, 1183–1193. https://doi.org/10.1016/S0092-8674(00)81462-2

Nicolay, B.N., Danielian, P.S., Kottakis, F., Jr, J.D.L., Sanidas, I., Miles, W.O., Dehnad, M., Tschöp, K., Gierut, J.J., Manning, A.L., Morris, R., Haigis, K., Bardeesy, N., Lees, J. a, Haas, W., Dyson, N.J., 2015. Proteomic analysis of pRb loss highlights a signature of decreased mitochondrial oxidative phosphorylation 1875–1889. https://doi.org/10.1101/gad.264127.115.4

Nicolay, B.N., Dyson, N.J., 2013. The multiple connections between pRB and cell metabolism. Curr. Opin. Cell Biol. 25, 735–740. https://doi.org/10.1016/j.ceb.2013.07.012

Nicolay, B.N., Gameiro, P.A., Tschöp, K., Korenjak, M., Heilmann, A.M., Asara, J.M., Stephanopoulos, G., Iliopoulos, O., Dyson, N.J., 2013. Loss of RBF1 changes glutamine catabolism. Genes Dev. 27, 182–96. https://doi.org/10.1101/gad.206227.112

Pasco, M.Y., Léopold, P., 2012. High sugar-induced insulin resistance in Drosophila relies on the Lipocalin Neural Lazarillo. PLoS One 7, 1–8. https://doi.org/10.1371/journal.pone.0036583

Pastor-Pareja, J.C., Xu, T., 2011. Shaping Cells and Organs in Drosophila by Opposing Roles of Fat Body-Secreted Collagen IV and Perlecan. Dev. Cell 21, 245–256. https://doi.org/10.1016/j.devcel.2011.06.026

Reynolds, M.R., Lane, A.N., Robertson, B., Kemp, S., Liu, Y., Hill, B.G., Dean, D.C., Clem, B.F., 2014. Control of glutamine metabolism by the tumor suppressor Rb. Oncogene 33, 556–566. https://doi.org/10.1038/onc.2012.635

Royzman, I., Whittaker, A.J., Orr-Weaver, T.L., 1997. Mutations in Drosophila DP and E2F distinguish G1-S progression from an associated transcriptional program. Genes Dev. 11, 1999–2011. https://doi.org/10.1101/gad.11.15.1999

Tennessen, J.M., Barry, W.E., Cox, J., Thummel, C.S., 2014. Methods for studying metabolism in Drosophila. Methods 68, 105–115. https://doi.org/10.1016/j.ymeth.2014.02.034

Ting, L., Rad, R., Gygi, S.P., Haas, W., 2011. MS3 eliminates ratio distortion in isobaric multiplexed quantitative proteomics. Nat. Methods 8, 937–940. https://doi.org/10.1038/nmeth.1714

Van Den Heuvel, S., Dyson, N.J., 2008. Conserved functions of the pRB and E2F families. Nat. Rev. Mol. Cell Biol. 9, 713–724. https://doi.org/10.1038/nrm2469

Watson, K.L., Johnson, T.K., Denell, R.E., 1991. Lethal(1)Aberrant Immune Response Mutations Leading to Melanotic Tumor Formation in Drosophila melanogaster. Dev. Genet. 12, 173– 187.

Yasugi, T., Yamada, T., Nishimura, T., 2017. Adaptation to dietary conditions by trehalose metabolism in Drosophila. Sci. Rep. 7, 2–10. https://doi.org/10.1038/s41598-017-01754-9

Zappia, M.P., Frolov, M. V, 2016. E2F function in muscle growth is necessary and sufficient for viability in Drosophila. Nat Commun 7:10509, 1–16. https://doi.org/10.1038/ncomms10509

Zhao, X., Karpac, J., 2017. Muscle Directs Diurnal Energy Homeostasis through a Myokine-Dependent Hormone Module in Drosophila. Curr. Biol. 1–15. https://doi.org/10.1016/j.cub.2017.06.004

